# Inter-subject pattern analysis: a straightforward and powerful scheme for group-level MVPA

**DOI:** 10.1101/587899

**Authors:** Qi Wang, Bastien Cagna, Thierry Chaminade, Sylvain Takerkart

## Abstract

Multivariate pattern analysis (MVPA) has become vastly popular for analyzing functional neuroimaging data. At the group level, two main strategies are used in the literature. The standard one is hierarchical, combining the outcomes of within-subject decoding results in a second-level analysis. The alternative one, inter-subject pattern analysis, directly works at the group-level by using, e.g, a leave-one-subject-out cross-validation. This study provides a thorough comparison of these two group-level decoding schemes, using both a large number of artificial datasets where the size of the multivariate effect and the amount of inter-individual variability are parametrically controlled, as well as two real fMRI datasets comprising respectively 15 and 39 subjects. We show that these two strategies uncover distinct significant regions with partial overlap, and that inter-subject pattern analysis is able to detect smaller effects and to facilitate the interpretation. The core source code and data are openly available, allowing to fully reproduce most of these results.

## 1. Introduction

Over the past decade, multi-voxel pattern analysis (MVPA, Haxby et al. (2014)) has become a very popular tool to extract knowledge from functional neuroimaging data. The advent of MVPA has offered new opportunities to examine neural coding at the macroscopic level, by making explicitly usable the information that lies in the differential modulations of brain activation across multiple locations – i.e multiple sensors for EEG and MEG, or multiple voxels for functional MRI (fMRI). Performing an MVP analysis commonly consists in *decoding* the multivariate information contained in functional patterns using a classifier that aims to guess the nature of the cognitive task performed by the participant when a given functional pattern was recorded. The decoding performance is consequently used to measure the ability of the classifier to distinguish patterns associated with the different tasks included in the paradigm. It provides an estimate of the *quantity of information* encoded in these patterns, which can then be exploited to localize such informative patterns and/or to gain insights on the underlying cognitive processes involved in these tasks.

This decoding performance is classically estimated separately on each of the participants. At the group level, these within-subject measurements are then combined – often using a *t*-test – to provide population-based inference, similarly to what is done in the standard hierarchical approach used in activation studies. Despite several criticisms of this group-level strategy that have been raised, namely on the nature of the statistical distribution of classification accuracies Olivetti et al. (2012), on the non-directional nature of the identified group-information Gilron et al. (2017) or on the fact that the results can be biased by confounds Todd et al. (2013), this hierarchical strategy remains widely used.

An alternative strategy directly works at the group-level by exploiting data from all available individuals in a single analysis. In this case, the decoding performance is assessed on data from new participants, i.e participants who did not provide data for the training of the classifier (see e.g Takerkart et al. (2014); Helfinstein et al. (2014); Jiang et al. (2016); Kim et al. (2016); Izuma et al. (2017); Etzel et al. (2016)), ensuring that the *information nature* is consistent across all individuals of the population that was sampled for the experiment. This strategy takes several denominations in the literature such as across-, between- or inter-subject classification or subject-transfer decoding. We hereafter retain the name inter-subject pattern analysis (ISPA).

In this paper, we describe a comparison of the results provided by these two classifier-based group-level MVPA strategies, which, to the best of our knowledge, is the first of its kind. This experimental study was carefully designed to exclusively focus on the differences induced by the within- vs. inter-subject nature of the decoding, i.e by making all other steps of the analysis workflow strictly identical. We provide results for both two real fMRI datasets and a large number of artificial datasets where the characteristics of the data are parametrically controlled. This allows us to demonstrate that these strategies offer different detection power, with a clear advantage for the inter-subject scheme, but furthermore that they can provide results of different nature, for which we put forward a potential explanation supported by the results of our simulations of the artificial data. The paper is organized as follows. Section. 2 describes our methodology, including our MVPA analysis pipeline for the two group-level strategies, as well as a description of the real datasets and the generative model of the artificial datasets. Section 3 includes the comparison of the results obtained with both strategies on these data, both in a qualitative and quantitative way. Finally, in Section 4, we discuss the practical consequences of our results and formulate recommendations for group-level MVPA.

## 2. Methods

### 2.1. Group-level Within-Subject Pattern Analysis (G-WSPA)

Since the seminal work of Haxby et al. (2001) that marked the advent of multivariate pattern analysis, most MVPA studies have relied on a within-subject decoding paradigm. For a given subject, the data is split between a training and a test set, a classifier is learnt on the training set and its generalization performance – usually measured as the classification accuracy – is assessed on the test set. If this accuracy turns out to be above chance level, it means that the algorithm has identified a combination of features in the data that distinguishes the functional patterns associated with the different experimental conditions. Said otherwise, this demonstrates that the input patterns contain information about the cognitive processes recruited when this subject performs the different tasks that have been decoded. The decoding accuracy can then be used as an estimate of the *amount* of available information – the higher accuracy, the more distinguishable the patterns, the larger the amount of information.

The group-level extension of this procedure consists in evaluating whether such information is present throughout the population being studied, which has led to the term *information-based brain mapping* Kriegeskorte et al. (2006). For this, a second level statistical analysis is conducted, for instance to test whether the average classification accuracy (or any other relevant summary statistic measured at the single-subject level), computed over the group of participants, is significantly above chance level. This can be done using a variety of approaches (see 2.5 for references). This hierarchical scheme is the one that is most commonly used in the literature. We denote it as Group-level Within-Subject Pattern Analysis (G-WSPA) in the rest of the present paper and illustrate on Fig. 1.

**Figure 1:**
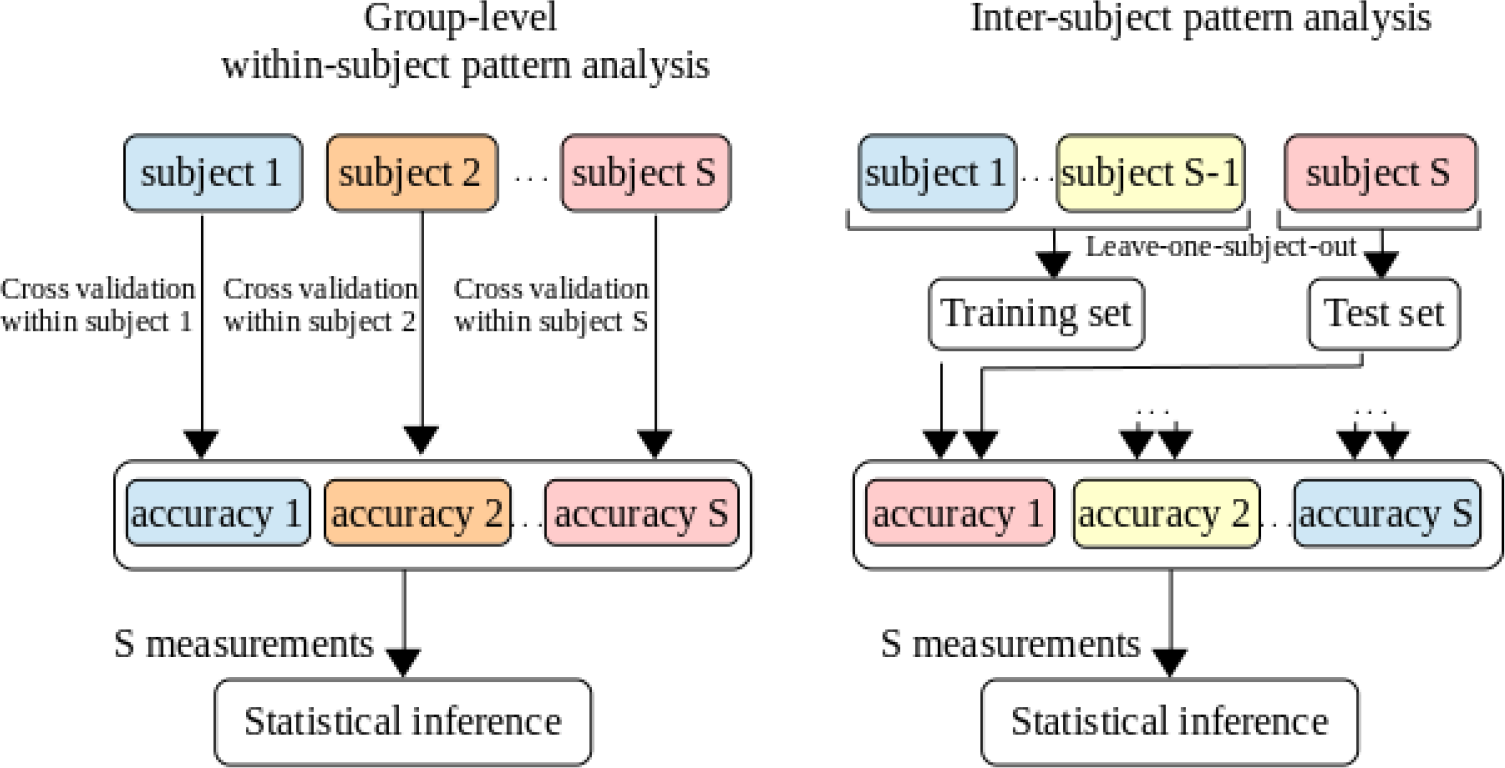
Illustration of the two approaches available to perform classifier-based group-level MVPA. Left: group-level within-subject pattern analysis (G-WSPA). Right: inter-subject pattern analysis (ISPA). Note that since we use a leave-one-subject-out cross-validation for ISPA, the two approaches yield the same number of measurements (equal to the number of subjects *S*), which allows for an unbiased comparison using the same statistical inference method.

### 2.2. Inter-Subject Pattern Analysis (ISPA)

Besides the hierarchical G-WSPA solution, another classifier-based framework exists to evaluate multivariate effects at the group level. Considering the data from all available individuals, one can train a classifier on data from a set of subjects – the training subjects – and evaluate its generalization capability on data from the others – the test subjects. One then use a cross-validation scheme that shuffles the subjects between the training and test sets, such as leave-one-subject-out or leave-n-subjects-out. In this setting, obtaining an average classification accuracy – this time across folds of the cross-validation – significantly above chance level means that a multivariate effect has been identified and is consistent across individuals, i.e that the multivariate information is of the same *nature* throughout the population. We denote this strategy as Inter-Subject Pattern Analysis (ISPA).

In this study, we use a leave-one-subject-out cross-validation in which the model accuracy is repeatedly computed on the data from the left-out subject. Even if other schemes might be preferable to multiply the number of measurements Varoquaux (2017), this choice was made to facilitate the comparison of the results obtained with ISPA and G-WSPA, as illustrated on Figure. 1.

### 2.3. Artificial data

The first type of data we use to compare G-WSPA and ISPA is created artificially. We generate a large number of datasets in order to conduct numerous experiments and obtain robust results. Each dataset is composed of 21 subjects (for ISPA: 20 for training, 1 for testing), with data points in two classes labeled in 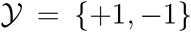, simulating a paradigm with two experimental conditions. For a given dataset, each subject *s* ∈ {1, 2, …, 21} provides 200 labeled observations, 100 per class. We note the *i*-th observation and corresponding class label 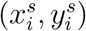, where 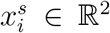 and 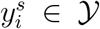. The pattern 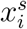 is created as

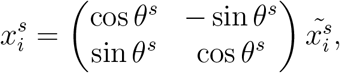

where

- 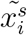 is randomly drawn from a 2D Gaussian distribution, respectively 𝓝(*C*^+^, Σ) and 𝓝(*C*^−^, Σ) if 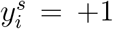 or 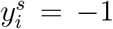, which are defined by their centers 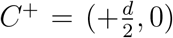 and 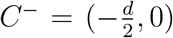, where *d* ∈ ℝ^+^ and their covariance matrix Σ, here fixed to 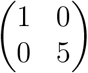 (see Supplementary Materials for results with other values of Σ);
- *θ*^*s*^ defines a rotation around the origin that is applied to all patterns of subjects *s*; the value of *θ*^*s*^ is randomly drawn from the Gaussian distribution 𝓝(0, Θ), where Θ defines the within-population variance.

Let 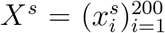 and 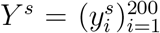 be the set of patterns and labels for subject *s*. A full dataset *D* is defined by

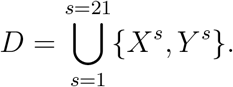

The characteristics of such a dataset are in fact governed by two parameters:

- *d*, which defines the distance between the point clouds of each of the two classes, i.e the multivariate effect size;
- Θ, which controls the amplitude of the rotation that can be applied to the data, separately for each subject: when Θ is small, all the *θ^s^* angles remain small, which means that the data of all subjects are similar; when Θ increases, the differences between subjects become larger; therefore, Θ quantifies the amount of inter-individual variability that exists within the group of 21 subjects for a given dataset.

Figures 2a and 2b illustrate the effect of each of these two parameters. Figure 2c shows different datasets generated with the same values of *d* and Θ.

**Figure 2:**
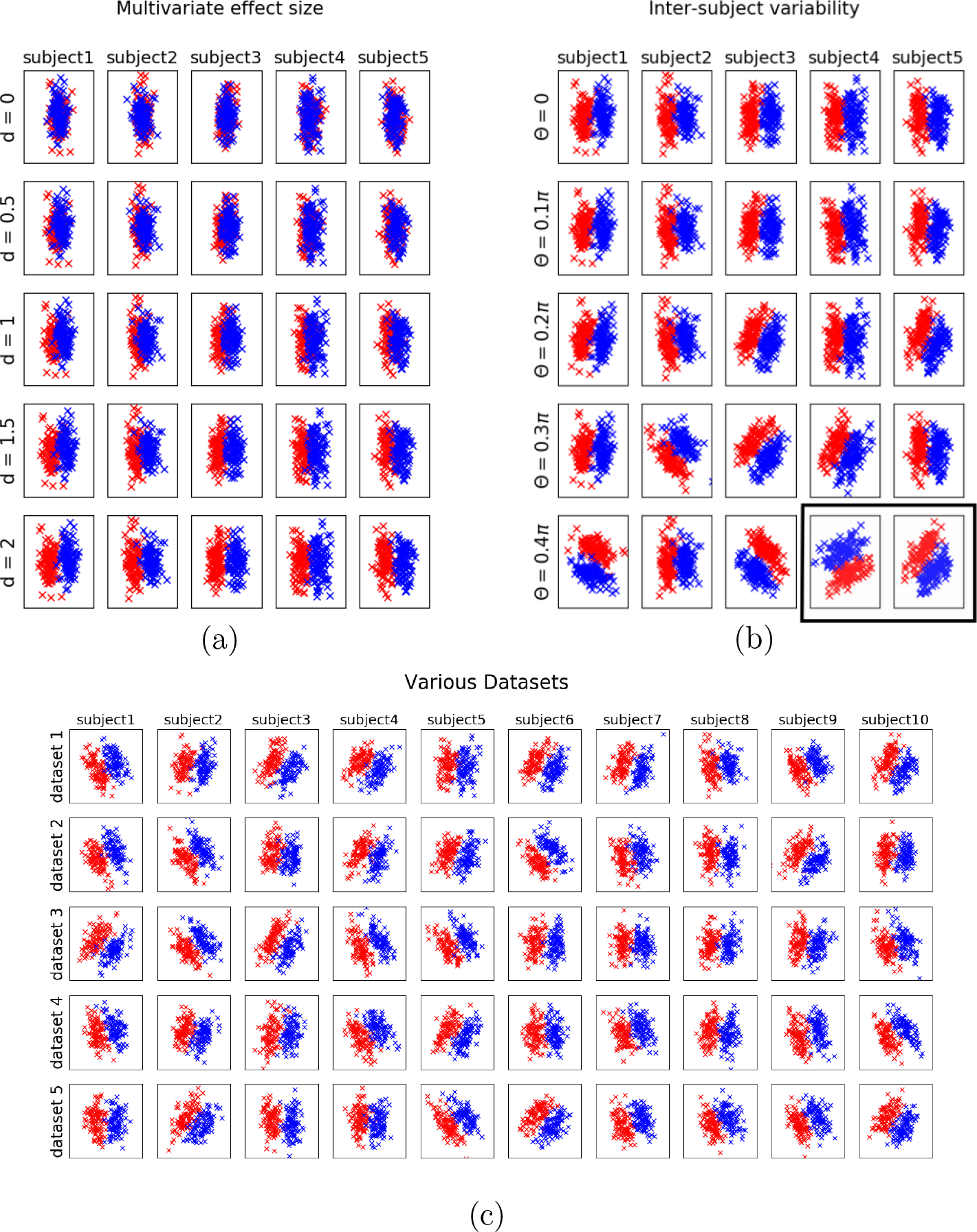
Illustration of the artificial datasets generated with the model described in 2.3. Each line is a subpart of a single dataset (5 subjects shown amongst 21 in (a) and (b), 10 subjects shown in (c)). The data points belonging to the class *y* = +1 and *y* = −1 are shown in blue and red, respectively. (a): influence of the *d* parameter (increasing effect size from top to bottom). (b): influence of the Θ parameter (increasing inter-individual variability from top to bottom). (c) five datasets obtained with the same values of the two parameters (*d* = 2 and Θ = 0.2*π*).

In our experiments we used 13 values for *d* and 11 values for Θ, *d* ∈ {0.1,0.12,0.14,0.16,0.18,0.2,0.22,0.24,0.26,0.28,0.3,0.4,0.6}, Θ ∈ {0.2*π*, 0.25*π*, 0.3*π*, 0.35*π*, 0.4*π*, 0.45*π*, 0.5*π*, 0.55*π*, 0.6*π*, 0.65*π*, 0.7*π*}, which gives 143 points in the two dimensional parameter space spanned by *d* and Θ. Note that by changing the value of Θ while keeping Σ constant, we control the relative amounts of within- and between-subject variance, which have been shown to be critical in group-level decoding situations Lindquist et al. (2017). For each pair (*d*, Θ), we generated 100 datasets. This yields 14300 datasets, each comprising 21 subjects and a total of 4200 data points. The code for generating these datasets (as well as performing the experiments detailed hereafter) is available online at the following URL: http://www.github.com/SylvainTakerkart/inter_subject_pattern_analysis.

### 2.4. fMRI data

We also used two real fMRI datasets that were acquired at the *Centre IRM-INT* in Marseille, France. For both experiments, participants provided written informed consent in agreement with the local guidelines of the South Mediterranean ethics committee.

In the first experiment (hereafter *Dataset1*), fifteen subjects participated to an investigation of the neural bases of cognitive control in the frontal lobe, largely reproducing the experimental procedure described in Koechlin and Jubault (2006). Participants lying supine in the MRI scanner were presented with audiovisual stimuli that carried complex information about the laterality of a button response, with the right or left thumb. Data was collected with a 3T Bruker Medspec 30/80 Avance scanner running ParaVision 3.0.2. Eight MRI acquisitions were performed. First, a field map using a double echo Flash sequence recorded distortions in the magnetic field. Six sessions with 60 trials each were recorded, each comprising 133 volumes (EPI sequence, isotropic resolution of 3 × 3 × 3*mm*, TE of 30*ms*, flip angle of 81.6°, field of view of 192 × 192*mm*, 36 interleaved ascending axial slices acquired within the TR of 2400*ms*) encompassing the whole brain parallel to the AC-PC plane. Finally, we acquired high-resolution T1-weighted anatomical images of each participant (MPRAGE sequence, isotropic voxels of 1 × 1 × 1*mm*, field of view of 256 × 256 × 180*mm*, *TR* = 9.4*ms*, *TE* = 4.424*ms*).

In the second experiment (*Dataset2*), thirty-nine subjects were scanned using a *voice localizer* paradigm, adapted from the one analyzed in Pernet et al. (2015). While in the scanner, the participants were asked to close their eyes while passively listening to a set of 144 audio stimuli, half of them being voice sounds, the other half being non-vocal. Most of the stimuli were taken from a database created for a previous study Capilla et al. (2012), while the others were extracted from copyright-free online databases. The paradigm was event-related, with inter-stimulus intervals randomly chosen between 4 and 5 seconds. The images were acquired on a 3T Prisma MRI scanner (Siemens, Eerlangen, Germany) with a 64-channels head coil. A pair of phase-reversed spin echo images was first acquired to estimate a map of the magnetic field. Then, a multi-band gradient echo-planar imaging (EPI) sequence with a factor of 5 was used to cover the whole brain and cerebellum with 60 slices during the TR of 955 ms, with an isotropic resolution of 2 × 2 × 2*mm*, a TE of 35.2 ms, a flip angle of 56 degrees and a field of view of 200 × 200*mm* for each slice. A total of 792 volumes were acquired in a single run of 12 minutes and 36 seconds. Then, a high resolution 3D T1 image was acquired for each subject (isotropic voxel size 0.8*mm*^3^, *TR* = 2400*ms*, *TE* = 2.28*ms*, field of view of 256 *×* 256 *×* 204.8*mm*). *Dataset2* is part of the InterTVA data set Aglieri et al. (2019), which is fully available online ^1^.

The two datasets were processed using the same sets of operations. The pre-processing steps were performed in SPM12^2^. They included co-registration of the EPIs with the T1 anatomical image, correction of the image distortions using the field maps, motion correction of the EPIs, construction of a population-specific anatomical template using the DARTEL method, transformation of the DARTEL template into MNI space and warping of the EPIs into this template space. Then, a general linear model was set up with one regressor per trial, as well as other regressors of non interest such as motion parameters. The estimation of the parameters of this model yielded a set of beta maps that was each associated with a given experimental trial. The beta values contained in these maps allowed constructing the vectors that serves as inputs to the decoders. We obtained 360 and 144 beta maps per subject for *Dataset1* and *Dataset2* respectively. No spatial smoothing was applied on these data for the results presented below (the results obtained with smoothing are provided as Supplementary Materials).

For these real fMRI datasets, we performed a searchlight decoding analysis Kriegeskorte et al. (2006), which allows to map local multivariate effects by sliding a spherical window throughout the whole brain and performing independent decoding analyses within each sphere. For our experiments, we exploited the searchlight implementation available in nilearn^3^ to allow obtaining the single-fold accuracy maps necessary to perform inference. For *Dataset1*, the decoding task was to guess whether the participant had used his left vs. right thumb to answer during the trial corresponding to the activation pattern provided to classifier. For G-WSPA, the within-subject cross-validation followed a leave-two-sessions-out scheme. For *Dataset2*, the binary classification task consisted in deciphering whether the sound presented to the participant was vocal or non-vocal. For G-WSPA, because a single session was available, we used an 8-fold cross-validation. Finally, all experiments were repeated with five different values of the searchlight radius (*r* ∈ {4*mm*, 6*mm*, 8*mm*, 10*mm*, 12*mm*}).

### 2.5. Statistical inference and performance evaluation

In order to perform statistical inference at the group level, the common practice is to use a *t-*test on the decoding accuracies. Such a test assesses whether the null hypothesis of a chance-level average accuracy can be rejected, which would reveal the existence of a multivariate difference between conditions at the group level. Note that as detailed in 2.1 and 2.2, even if the same statistical test is applied for both strategies, a rejection of the null hypothesis reflects different underlying phenomenons: for G-WSPA, it means that the *quantity* of information distinguishing the experimental conditions is on average non null throughout the population, whereas for ISPA, it further implies that the *nature* of the multivariate information is consistent across individuals.

However, it is now well established that several properties necessary for such parametric test to be valid are not met in this context (see e.g Stelzer et al. (2013)), such as the fact that the accuracy of a classifier does not follow a gaussian distribution. In order to overcome this limitation, we used a non parametric permutation test Nichols and Holmes (2002) to assess the significance of the measured *t* statistic, which allows revealing whether the decoding accuracy is significantly greater than chance at the group level, for all the experiments conducted in the present study. While other more sophisticated alternatives have been proposed in the literature (see e.g Olivetti et al. (2012); Brodersen et al. (2013); Stelzer et al. (2013); Etzel (2015); Allefeld et al. (2016)), the implementation of this procedure is straightforward, and it allows comparing the results given by G-WSPA and ISPA in a fair manner, which is the objective of the present study. In practice, for the real fMRI experiments, we used the implementation offered in the SnPM toolbox^4^ to analyse the within-subject (for G-WSPA) or the single-fold (for the inter-subject cross-validation of ISPA) accuracy maps, with 1000 permutations and a threshold classically chosen at *p <* 0.05 (FWE corrected). For the simulations that used the artificial datasets, we used an in-house implementation of the equivalent permutation test (also available in our open source code; see 2.3), with 1000 permutations and a threshold at *p <* 0.05. Critically, it should be noted that the same number of samples were available for this statistical procedure when using G-WSPA or ISPA, as shown on Figure. 1.

In order to compare the group-level decoding results provided by GWSPA and ISPA, we use the following set of metrics. For the artificial data, we generated 100 datasets at each of the 143 points of the two dimensional parameter space spanned by the two parameters *d* and Θ. For each of these datasets, we estimate the probability *p* to reject the null hypothesis of no group-level decoding. We then simply count the number of datasets for which this null hypothesis can be rejected, using the *p* < 0.05 threshold, which we respectively denote *N*_*G*_ and *N*_*I*_ for G-WSPA and ISPA. For the experiments conducted on the two real fMRI datasets, we examine the thresholded statistical map obtained for each experiment. We then compare the maps obtained by G-WSPA and ISPA by computing the size and maximum statistic of each cluster, as well as quantitatively assessing their extent and localization by measuring how they overlap.

## 3. Results

In this section, we present the results obtained when comparing G-WSPA and ISPA on both the artificial and real fMRI datasets. With the artificial datasets, our focus is on the characterization of the space spanned by the two parameters that control the characteristics of the data. On the real datasets, we quantitatively and qualitatively assess the statistic maps produced by these two strategies, examine the influence of the size of the searchlight beam, and try to relate these results to the one obtained on the artificial data.

### 3.1. Results for artificial datasets

The results of the application of G-WSPA and ISPA on the 14300 datasets that were artificially created are summarized in Tables S1 and S2. In order to facilitate grasping the results on this very large number of datasets, we proceed in two steps. First, we represent the two-dimensional parameter space spanned by *d* and Θ as a table where the columns and lines respectively represent a given value for these two parameters. The values (denoted as *N*_*G*_ for G-WSPA and *N*_*I*_ for ISPA) in these tables are the number of datasets (out of the 100 datasets available for each cell) for which a significant group level decoding accuracy (*p* < 0.05, permutation test) is obtained with G-WSPA (Table S1) or ISPA (Table S2). Secondly, we arbitrarily threshold these dataset counts, coloring only the cells where at least half of the datasets yield significant results (in green for G-WSPA when *N*_*G*_ ⩾ 50 and red for ISPA when *N*_*I*_ ⩾ 50).

**Table S1:** Number of datasets *N*_*G*_ (out of 100) for which G-WSPA provides significant group decoding (in green: cells where *N*_*G*_ ≥ 50)

| effect size<br>variability | 0.1 | 0.12 | 0.14 | 0.16 | 0.18 | 0.2 | 0.22 | 0.24 | 0.26 | 0.28 | 0.3 | 0.4 | 0.6 |
| --- | --- | --- | --- | --- | --- | --- | --- | --- | --- | --- | --- | --- | --- |
| $0.7\pi$ | 7 | 11 | 15 | 22 | 27 | 36 | 40 | 44 | 53 | 59 | 68 | 96 | 100 |
| $0.65\pi$ | 9 | 14 | 16 | 23 | 29 | 36 | 39 | 45 | 55 | 63 | 70 | 98 | 100 |
| $0.6\pi$ | 13 | 16 | 17 | 25 | 27 | 35 | 44 | 53 | 58 | 71 | 76 | 95 | 100 |
| $0.55\pi$ | 10 | 14 | 17 | 21 | 25 | 32 | 37 | 46 | 50 | 61 | 76 | 96 | 100 |
| $0.5\pi$ | 7 | 8 | 15 | 17 | 22 | 30 | 36 | 46 | 55 | 59 | 64 | 94 | 100 |
| $0.45\pi$ | 4 | 7 | 13 | 14 | 22 | 32 | 39 | 46 | 51 | 62 | 69 | 96 | 100 |
| $0.4\pi$ | 8 | 11 | 18 | 22 | 26 | 31 | 40 | 49 | 54 | 63 | 72 | 97 | 100 |
| $0.35\pi$ | 4 | 7 | 15 | 24 | 34 | 37 | 46 | 51 | 61 | 61 | 68 | 96 | 100 |
| $0.3\pi$ | 10 | 14 | 24 | 26 | 31 | 40 | 46 | 57 | 58 | 71 | 78 | 95 | 100 |
| $0.25\pi$ | 9 | 13 | 20 | 25 | 32 | 43 | 45 | 53 | 60 | 62 | 69 | 94 | 100 |
| $0.2\pi$ | 9 | 12 | 15 | 19 | 28 | 34 | 39 | 50 | 59 | 67 | 75 | 95 | 100 |

**Table S2:**
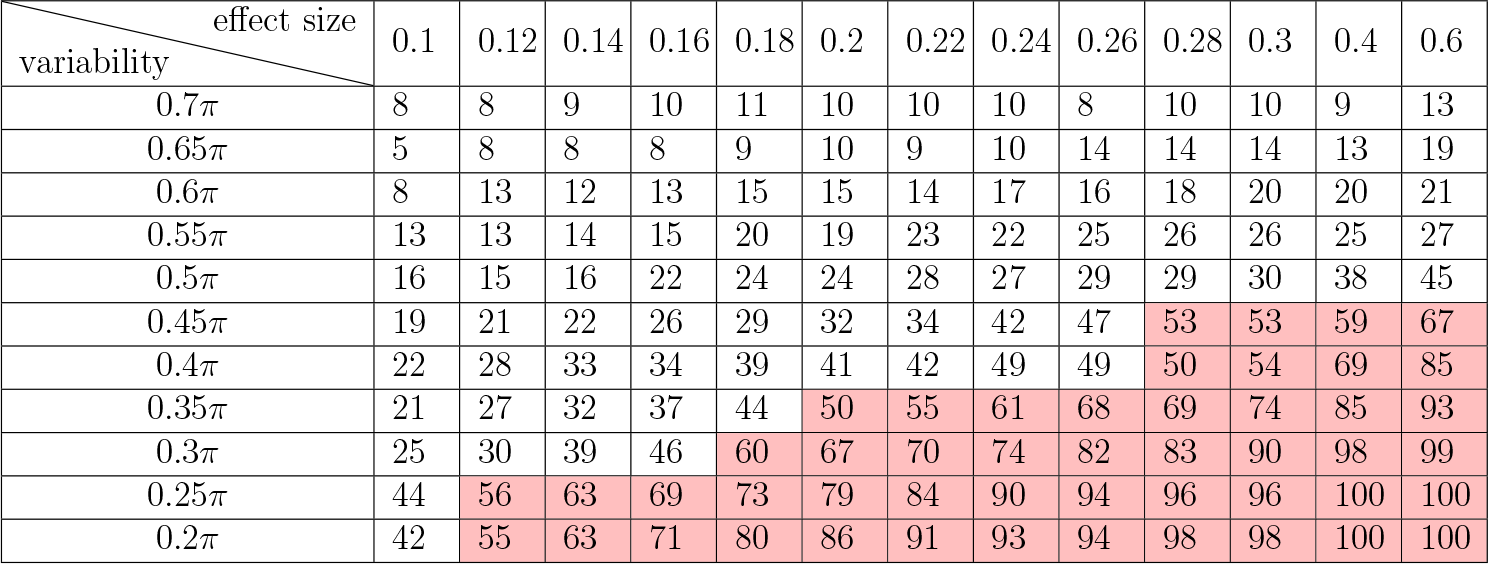
Number of datasets *N*_*I*_ (out of 100) for which ISPA provides significant group decoding (in red: cells where *N*_*I*_ ≥ 50)

**Table S3:**
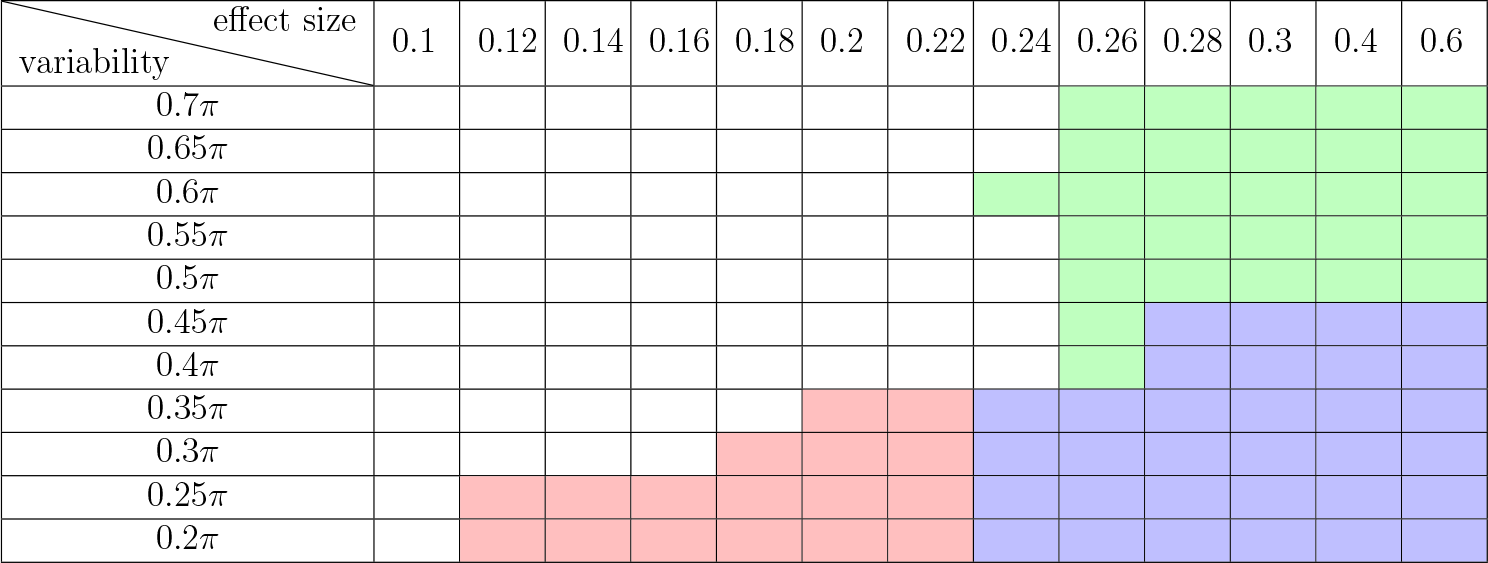
Visual comparison of G-WSPA vs ISPA (in blue: cells where both *N*_*G*_ ≥ 50 and *N*_*I*_ ≥ 50; in green: cells where *N*_*G*_ ≥ 50 and *N*_*I*_ < 50; in red: cells where *N*_*G*_ < 50 and *N*_*I*_ ≥ 50).

Table S1 shows that i) in any given column, the number *N*_*G*_ of datasets for which G-WSPA detects a significant effect seem to vary weakly, suggesting that the influence of the amount of inter-individual variability (i.e the value of Θ) is fairly small on the outcome of the G-WSPA strategy; ii) in any given line, *N*_*G*_ decreases with *d*, from a value of 100 (i.e all datasets) for the largest tested effect size (*d* = 0.3). This produces the rectangle-like area visible in green on Table S1. Table S2 shows that the results of ISPA are more complex to explain, with a clear influence of both *d* and Θ. When the inter-individual variability is low, ISPA can detect significant effects even with very small effect sizes. When the variability increases, the detection power of ISPA decreases – i.e for a given effect size, the number of datasets for which ISPA yields a significant result decreases. Therefore the detection power of ISPA is determined by a trade-off between *d* and Θ, which produces the triangle-like area visible in red on Table S2. These observations are confirmed by the results of 2-way analyses of variance performed on the content of each of these two tables, i.e in which we used two factors, *d* and Θ to try to model the number of data sets for which we obtained significant group-decoding. The effects of *d* and Θ are significant in both tables, but it is the effect of Θ in Table S1 that is, by far, the weakest: 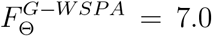, *p* = 1.2*e*^−8^, compared to 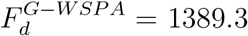, *p* < 1*e*^−121^; 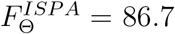, *p* < 1*e*^−49^, and 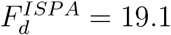, *p* < 1*e*^−21^.

In order to easily depict the compared behaviors of G-WSPA and ISPA, we *overlapped* the results of the two strategies into Table S3. In this third table, the blue cells indicate that *N*_*G*_ ⩾ 50 and *N*_*I*_ ⩾ 50 (i.e that both G-WSPA and ISPA produce significant results in more than half of the 100 datasets), while the green and red regions contain cells where it is the case only for G-WSPA or ISPA respectively (i.e *N*_*G*_ ⩾ 50 and *N*_*I*_ < 50 in green cells; *N*_*I*_ ⩾ 50 and *N*_*G*_ < 50 in red cells). We observe a large blue region in which both strategies provide concordant results, for the largest values of the effect size *d* and with a moderately low amount of inter-individual variability. Interestingly, the green and red regions, where one strategy detects a group-level effect while the other does not, also take an important area in the portion of the parameter space that was browsed by our experiments, which means that the two strategies can disagree. G-WSPA can provide a positive detection when the inter-individual variability is very large, while ISPA cannot (green region). But ISPA is the only strategy that offers a positive detection for very small effect sizes, requiring a moderate inter-individual variability (red region).

### 3.2. Results for fMRI datasets

#### 3.2.1. Qualitative observations

The searchlight decoding analyses performed at the group level were all able to detect clusters of voxels where the decoding performance was significantly above chance level (*p <* 0.05, FWE-corrected using permutation tests) with both G-WSPA and ISPA, for *Dataset1* and *Dataset2* and with all sizes of the spherical searchlight. The detected clusters were overall consistent across values of the searchlight radius, with an increasing size of each cluster when the radius increases. In *Dataset1*, both strategies uncovered two large significant clusters located symmetrically in the left and right motor cortex. Additionally, ISPA was able to detect other significant regions located bilaterally in the dorsal part of the cerebellum and the parietal operculum, as well as a medial cluster in the supplementary motor area (note that some of these smaller clusters also become significant with G-WSPA with the larger searchlight radii). In *Dataset2*, both G-WSPA and ISPA yielded two large significant clusters in the temporal lobe in the left and right hemispheres, which include the primary auditory cortex as well as higher level auditory regions. Figure. 3 provides a representative illustration of these results, for a radius of 6*mm*.

**Figure 3:**
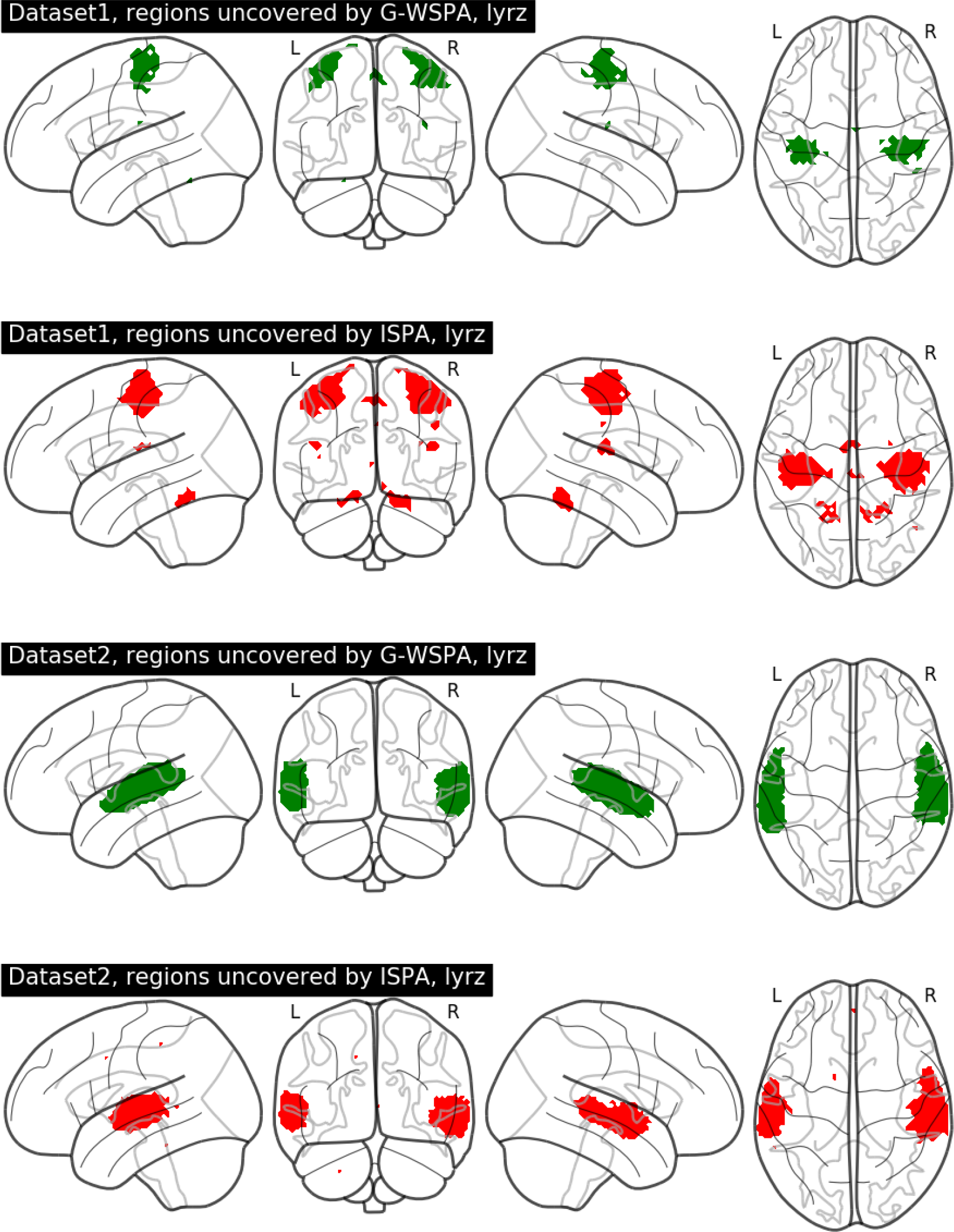
Illustration of the results of the group-level searchlight decoding analysis for a 6*mm* radius. Top two rows: *Dataset1*; bottom two rows: *Dataset2*. Brain regions found significant using G-WSPA and ISPA are respectively depicted in green and red.

#### 3.2.2. Quantitative evaluation

Our quantitative evaluation focuses on the two largest clusters uncovered in each dataset, i.e the ones in the motor cortex for *Dataset1* and the ones in the temporal lobe for *Dataset2*. We first examine the size of these clusters, separately for each hemisphere and each of the five values of the searchlight radius. The results are displayed in the left column of Figure. 4. In almost all cases, the size of the significant clusters increased with the searchlight radius (left column). Moreover, the cluster located in the right hemisphere is consistently larger than the one on the left. In *Dataset1*, the cluster detected by ISPA is larger than the one detected by G-WSPA, regardless of the hemisphere, while in *Dataset2*, it is G-WSPA that yields larger clusters (except for a 4*mm* radius where the sizes are similar). Then, we study the peak value of the *t* statistic obtained in each cluster (right column of Figure. 4). In *Dataset1*, the peak *t* value is higher for ISPA than G-WSPA, for all values of the radius. In *Dataset2*, ISPA yields higher peak *t* values than G-WSPA for the searchlight radii smaller or equal than 8*mm*, and lower peak *t* values for the larger radius values.

**Figure 4:**
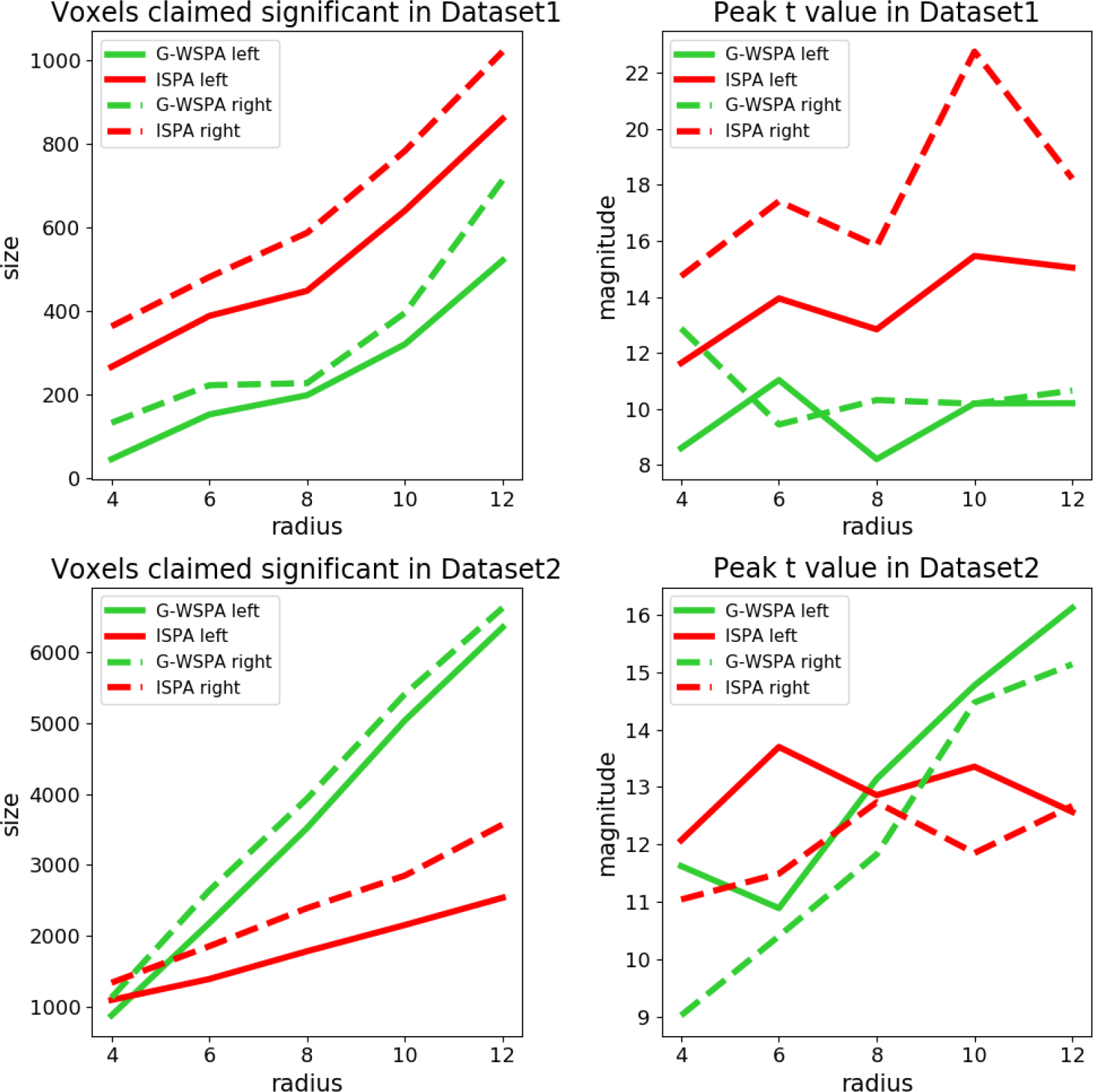
Quantitative evaluation of the results obtained on the real fMRI datasets for G-WSPA (green curves) and ISPA (red curves). Solid and dashed lines for the largest cluster respectively in the left and right hemispheres. Left column: size of the significant clusters. Right column: peak *t* statistic. Top vs. bottom row: results for *Dataset1* and *Dataset2* respectively.

Then, we quantify the amount of overlap between the clusters found by G-WSPA and ISPA, by splitting the voxels into three sub-regions: voxels uncovered only by ISPA, only by G-WSPA or by both strategies (overlap). Figure. 5 provides an illustration of these sub-regions, which shows that the overlap region (in blue) is located at the core of the detected clusters, while the voxels significant for only one strategy are located in the periphery; these peripheral voxels appear to be mostly detected by ISPA for *Dataset1* (red voxels) and by G-WSPA for *Dataset2* (green voxels). Figure. 6 shows the voxel counts in each sub-region, which confirms this visual inspection. Overall, the size of the sub-region found by the two strategies increases with the searchlight radius. The ISPA-only sub-region is larger in *Dataset1* than in *Dataset2*, representing between 38% and 83% of all significant voxels. Conversely in *Dataset2*, the G-WSPA-only sub-region is more important – with a percentage of all significant voxels comprised between 18% and 60%.

**Figure 5:**
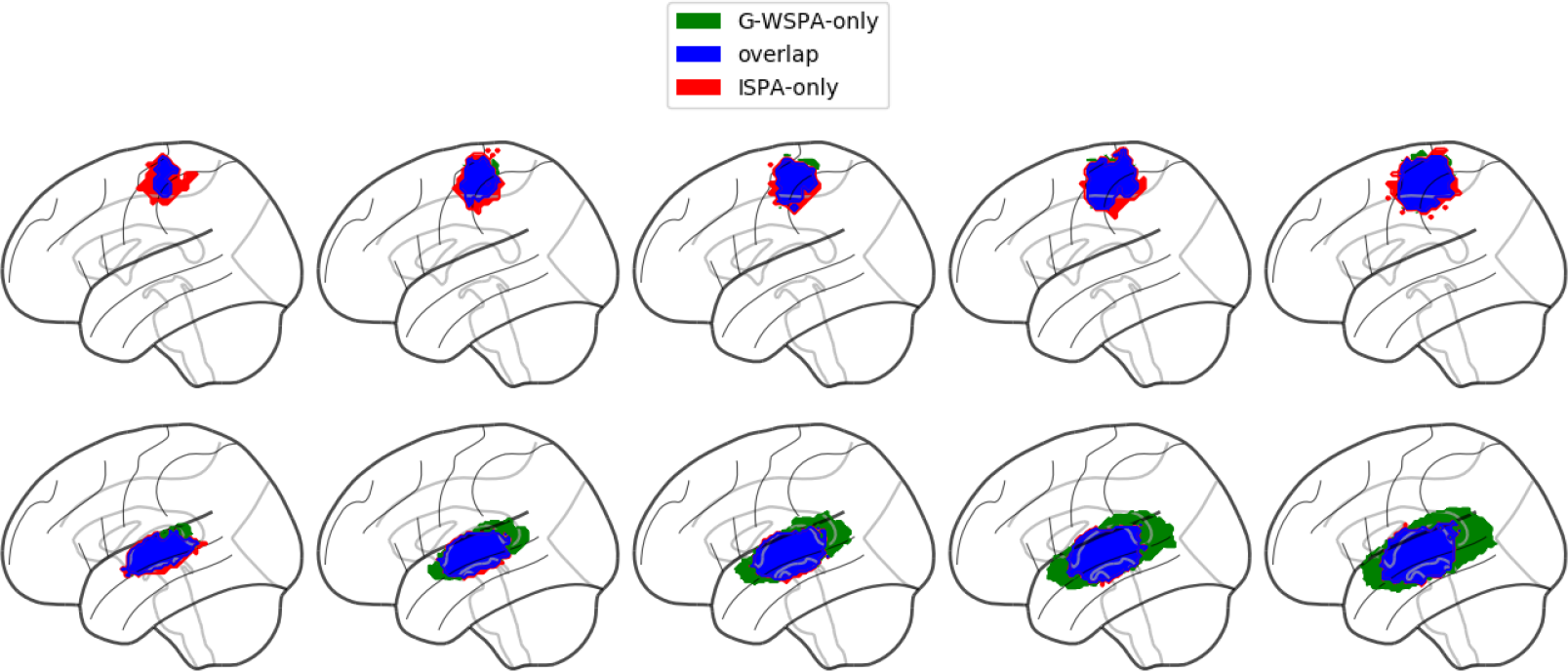
Comparison of the clusters detected by G-WSPA and ISPA for the different values of the searchlight radius, in *Dataset1* (top row) and *Dataset2* (bottom row).

**Figure 6:**
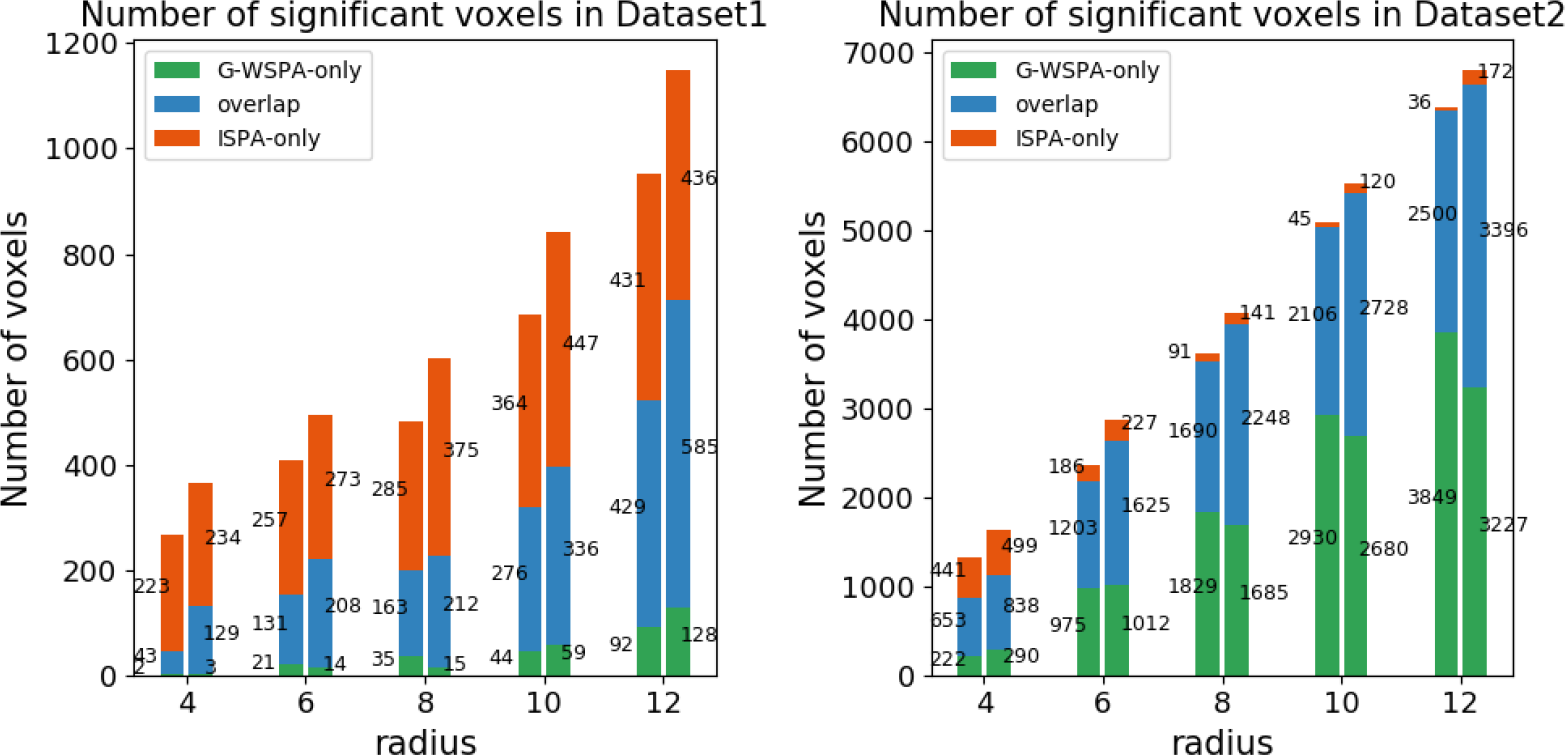
Comparison of the voxel counts detected by G-WSPA-only (green), ISPA-only (red) or both (blue) for the different values of the searchlight radius, in *Dataset1* (left) and *Dataset2* (right).

We also count the number of voxels in each sub-region for each hemisphere. Figure 6 shows that for both datasets, the clusters in the right hemisphere are larger than in the left hemisphere. For both hemispheres, in most cases the number of voxels in each sub-region increases as the search-light radius increases. However, in *Dataset1*, the number of voxels found only by G-WSPA is much smaller than that of overlap and ISPA-only sub-region with all five radius values. While in *Dataset2* the number of voxels in ISPA-only sub-region decreases for the four smallest values of the searchlight radius, and voxels only uncovered by ISPA are much fewer than those found only by G-WSPA.

## 4. Discussion

### 4.1. G-WSPA and ISPA provide different results

In this study, we have performed experiments on both real and artificial functional neuroimaging data in order to compare two group-level MVPA schemes that rely on classifier-based decoding analyses: the vastly used G-WSPA, and ISPA. Our results show that both strategies can offer equivalent results in some cases, i.e that they both detect significant group-level multi-variate effects in similar regions of the cortex for our two real fMRI datasets, and in parts of the two-dimensional parameter space browsed using our artificially generated datasets, but that their outcomes can also differ significantly. For instance, in *Dataset1*, ISPA was the only strategy that detected multivariate group-level effects in several regions such as the supplementary motor area, the bilateral parietal operculum and dorsal cerebellum, for most of the searchlight radii that we tested. Furthermore, when a cluster is detected by both strategies, it usually differs in its size, extent and/or precise location, resulting in partial overlap; in most cases, the areas of concordance between the two strategies appeared to be centrally located in the cluster, while the disagreements are located towards the periphery: in some areas, G-WSPA detects a group-level effect while ISPA does not, and inversely in other areas. Note that for *Dataset2*, our observations are consistent with what was reported in Gilron et al. (2017) with a different, yet comparable, framework of analysis, on a closely related data set.

Surprisingly, the peripheral behaviors were not consistent across the two real fMRI datasets: on *Dataset1*, ISPA only was able to detect effects on the periphery of the core region where both strategies were equally effective, while on *Dataset2*, it was G-WSPA which provided significant results on the periphery. The results of the experiments conducted on the artificial datasets can actually shed some light on these results, thanks to the clear dissociation that was observed in the two-dimensional parameter space browsed to control the properties of the data. Indeed, ISPA is the only strategy that allows detecting smaller multivariate effects when the inter-individual variability remains moderate, which is the case in the main clusters detected in *Dataset1* because they are located in the primary motor cortex, the *primary* nature of this region limiting the amount of inter-subject variability. On the opposite, the peripheral part of the main clusters detected in *Dataset2* are located anteriorly and posteriorly to the primary auditory cortex, towards higher-level auditory areas where the inter-individual variability is higher, a situation in which G-WSPA revealed more effective in the experiments conducted with our artificial data.

### 4.2. ISPA: larger training sets improve detection power

Our experiments revealed a very important feature offered by the ISPA strategy: its ability to detect smaller multivariate effects. On the one hand, this greater detection power was explicitly demonstrated through the simulations performed on artificial data, where the multivariate effect size was one of the two parameters that governed the generation of the data; we showed that with an equal amount of inter-individual variability, ISPA was able to detect effects as small as half of what can be detected by G-WSPA. Furthermore, on both real fMRI datasets, ISPA was able to detect significant voxels that were not detected using G-WSPA, in a large amount in *Dataset1*, and to a lesser extent in *Dataset2*. This detection power advantage is of great importance, since detecting weak distributed effects was one of the original motivations for the use of MVPA Haxby et al. (2014).

This greater detection power of ISPA is in fact the result of the larger size of the training set available: indeed, when the number of training examples is small, the performance of a model overall increases with the size of the training set, until an asymptote that is reached with large training sets – as encountered in computer vision problems where millions of images are available from e.g http://www.image-net.org. In the case of functional neuroimaging where an observation usually corresponds to an experimental trial, we usually have a few dozens to a few hundreds samples per subject, which clearly belongs to the *small sample size* regime, i.e very far from the asymptote. In this context, ISPA offers the advantage to multiply the number of training samples by a factor equal to the number of subjects in the training set, which is of great value. However, here, the increased sample size comes at the price of a larger heterogeneity in the training set, because of the large differences that can exist between data points recorded in different subjects. In the general case, such an added heterogeneity can represent an obstacle for learning if it is very important, but can also reveal beneficial if more moderate by increasing the diversity of the training set. The fact that we observe a higher detection power with ISPA than with G-WSPA suggests that we are in the latter situation.

### 4.3. ISPA offers straightforward interpretation

When using the ISPA strategy, obtaining a positive result means that the model has learnt an implicit rule from the data available in the training subjects that provides statistically significant generalization power on data from new subjects. Since a cross-validation of the type leave-one-subject-out or leave-n-subjects-out is performed on the available data to quantitatively assess such results, it allows to draw inference on the full population from which the group of participants was drawn, including individuals for which no data was available. As previously pointed out in Kragel et al. (2018), the interpretation that follows is straightforward: a significant effect detected with ISPA implies that some information has been identified to be consistent throughout the full population. In more details, this means that the modulations of the multivariate patterns according to the experimental conditions that were the object of the decoding analysis are consistent throughout the population, at least at the resolution offered by the modality used for the acquisition. This is the case for all voxels colored blue and red on Figure. 5.

In the case of a significant result detected by G-WSPA but not ISPA – i.e the green voxels on Figure. 5, the interpretation is not as straightforward. Such a result indeed implies that on average over all subjects of the population, there is information in the input patterns that can discriminate the different experimental conditions – since it is significant for G-WSPA, but that the nature of the discriminant information present in the input voxels differs across individuals – since it is not detected by ISPA. This could be caused by two phenomenons. First, it could mean that the underlying coding strategy is nonetheless invariant across individuals, but that the nature of the data or of the feature space used in this analysis does not allow to identify it as such. One would then need to acquire additional data using a different modality (Dubois et al. (2015)) or to transform the feature space (e.g using methods such as Haxby et al. (2011), Takerkart et al. (2014) or Fuchigami et al. (2018)) in order to attempt to make this invariance explicit. Secondly, it could also mean that the coding principle is simply intrinsically different across subjects (as might be the case for the two subjects framed at the bottom right of Figure 2b, in a dataset for which G-WSPA provides a positive detection), for instance because several strategies had been employed by different individuals to achieve the same task, or because each subject employs its own idiosyncratic neural code. In this second hypothesis, one could cast some doubt on the validity of tagging these voxels as significant at the group-level. This risk does not only apply to results obtained using searchlight decoding analyses, but also to analyses performed in pre-defined regions of interest, which are extremely common in the literature. Even if these limitations have been pointed out previously in the literature, as in e.g Todd et al. (2013), Allefeld et al. (2016) or Gilron et al. (2017), we feel that the community should tackle this question more firmly. This could start by defining what a group-level multivariate analysis should seek – a consistent *amount* or *nature* of the information, or by promoting the ISPA strategy which allows directly circumventing such potential risks.

### 4.4. ISPA: a computational perspective

From a practical point of view, one can ask two questions that are critical if one would like to promote the use of ISPA. First, what is the computational cost of ISPA and how does it compare to the one of G-WSPA? And secondly, is it easy to implement ISPA with the existing MVPA software packages?

In order to address the former question, we first compare the number of classifiers that need to be trained for a full group analysis. Using G-WSPA, we need to train *K* classifiers per subject, where *K* is the chosen number of within-subject cross-validation folds, so *KS* classifiers in total (where *S* is the number of available subjects). For ISPA, the number of cross-validation folds equals to *S* (for leave-one-subject-out), meaning we need to train a total of *S* classifiers. The training time of a classifier also depends on the number of training examples: it is linear for classifiers such as logistic regression (when using gradient-based optimizers Dreiseitl and Ohno-Machado (2002)), and quadratic for e.g support vector machines Bottou and Lin (2007). Assuming we have *n* examples per subject, the number of training examples available for each classifier is 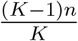 for G-WSPA and (*S* − 1)*n* for ISPA. Overall, with linear-time classifiers, the total complexity of a group-level decoding analysis amounts to *O*(*nKS*) for G-WSPA and *O*(*nS*^2^) for ISPA, which makes them almost equivalent if one assumes that *K* and *S* are of the same order of magnitude. With quadratic-time classifiers, the total complexity is *O*(*n*^2^*KS*) for G-WSPA and *O*(*n*^2^*S*^3^) for ISPA, which makes ISPA significantly more costly. We therefore advice to use linear-time classifiers such as logistic regression to perform ISPA analyses, particularly with searchlight decoding where the computational cost is further multiplied by the number of voxels.

In terms of software implementation, because within-subject analyses have been the standard since the advent of MVPA, all software packages provide well documented examples for such analyses which are the base tool for G-WSPA. Even if it is not the case for ISPA, it is easy to obtain an equivalent implementation because to perform inter-subject decoding, one simply need to i) have access to the data from all subjects, and ii) set up a leave-one-subject-out cross-validation, these two operations being available in all software packages. As an example, we provide the code to perform ISPA searchlight decoding from pre-processed data available online, which allows reproducing the results described in the present paper on *Dataset2*: http://www.github.com/SylvainTakerkart/inter_subject_pattern_analysis.

## 5. Conclusion

In this paper, we have compared two strategies that allow performing group-level decoding-based multivariate pattern analysis of task-based functional neuroimaging experiments: the first is the standard information-based method that aggregates within-subject decoding results and a second one that directly seeks to decode neural patterns at the group level in an inter-subject scheme. Both strategies revealed effective but they only provide partially concordant results. Inter-subject pattern analysis offers a higher detection power to detect weak distributed effects and facilitate the interpretation while the results provided by the information-based approach necessitate further investigation to raise potential ambiguities. Furthermore, because it allows identifying group-wise invariants from functional neuroimaging patterns, inter-subject pattern analysis is a tool of choice to identify neuromarkers Gabrieli et al. (2015) or brain signatures Kragel et al. (2018), making it a versatile scheme for population-wise multivariate analyses.

## Supporting information

Supplementary experiments

## Acknowledgments

This work was granted access to the HPC resources of Aix-Marseille Université financed by the project Equip@Meso (ANR-10-EQPX-29-01) of the French program *Investissements d’Avenir*. The acquisition of the data was made possible thanks to the infrastructure France Life Imaging (11-INBS-0006) of the French program *Investissements d’Avenir*, as well as specific grants from the *Templeton Foundation* (40463) and the *Agence Nationale de la Recherche* (ANR-15-CE23-0026).

1 https://openneuro.org/datasets/ds001771

2 https://www.fil.ion.ucl.ac.uk/spm/

3 http://nilearn.github.io/

4 http://warwick.ac.uk/snpm

