## Supplementary experiments for "Inter-subject pattern analysis: a straightforward and powerful scheme for group-level MVPA"

### Supplementary materials

---

We present here additional experiments that were conducted on the same fMRI datasets, as well as new artificial data that were created using the same generative model but different parameters.

#### 1. Influence of the number of subjects and number of samples per subject

**Aim**. Although the two real fMRI datasets used in our experiments have approximately the same total number of observations (5400 observations in *Dataset1*, 5616 observations in *Dataset2*), they differ in the number of subjects that were scanned and the number of trials per subject: *Dataset1* includes 360 trials for each of the 15 subjects, while *Dataset2* offers 144 trials for 39 subjects. We here attempt to investigate whether this could explain some of the differences we observed in the results of G-WSPA and ISPA on these two datasets, using new artificial datasets.

**Experiments**. We therefore repeat the same set of experiments as in the paper, using new sets of 14300 datasets generated to maintain the size of the training set constant and modulating the ratio between the number of subjects $S$ and the number of samples per subject $N$. We used the following values for the couple $(S, N)$:
$(S, N) \in \{(9, 500), (11, 400), (17, 250), (21, 200), (51, 80), (101, 40), (201, 20), (401, 10)\}$, which allows maintaining the size of the group-level dataset approximately constant (and more particularly, the size of the ISPA training set is exactly constant at $(S - 1) \times N = 4000$).

**Results**. We present below the results we obtained, under the same summarized form as in Table S3 of the paper. Tables 1 to 8 represent the two-dimensional parameter space spanned by $d$ and $\Theta$, for various values of the $(S, N)$ couple. The cells colored in blue are those where G-WSPA and ISPA yielded significant group-level decoding for 50 or more datasets out of the 100, whereas in the green cells it is the case only for G-WSPA and in the red ones it is the case only for ISPA.

The shape of the region where G-WSPA yields 50 or more significant detections, corresponding here to the green and blue cells, remains approximately rectangular for all values of $(S, N)$. But when the number of subject $S$ increases, it is shifted to the right of the table, i.e G-WSPA is effective only for larger effect sizes. This is a direct consequence of the decreasing value of $N$, which is critical for within-subject decoding. For ISPA, the shape and location of the colored region (red and blue cells) where it is effective remains fairly constant, showing that ISPA is not or weakly affected by the ratio $N/S$ when the size of the training set is constant.

We now try to address the question of whether the difference in the $N/S$ ratio between *Dataset1* and *Dataset2* can explain the fact that the most peripheral regions of the main clusters are detected by ISPA for *Dataset1* and G-WSPA for *Dataset2*, in the light of the present results where we vary $N/S$ in artificial datasets. The $N/S$ ratio for *Dataset1* is $360/15 = 24$, while for *Dataset2*, it is $144/39 = 3.7$. We reformulate the previous question as follows: if the $N/S$ ratio of *Dataset1* were smaller (i.e closer to the one of *Dataset2*), could the peripheral voxels – which are red on Fig.5 of the paper, i.e are detected by ISPA – *become* green, i.e be detected only by G-WSPA? For this, consider a red cell in Table 3 (for which the $N/S$ ratio is the closest to the one of *Dataset1*): can it become green when the ratio $N/S$ decreases? We clearly see that it is not possible, i.e that all red cells in Table 3 are also red (in almost all cases) in Tables 4 to 8 where $N/S$ is smaller. We then ask the opposite question: if the $N/S$ ratio of *Dataset2* were larger (i.e closer to the one of *Dataset1*), could the peripheral voxels – which are green on Fig.5 of the paper, i.e are detected by G-WSPA – *become* red, i.e be detected only by ISPA? For this, consider a green cell in Table 5 (for which the $N/S$ ratio is the closest to the one of *Dataset2*): can it become red when the ratio $N/S$ increases? We clearly see that it is not possible, i.e that all green cells in Table 5 are also green (in almost all cases) in Tables 1 to 4 where the $N/S$ ratio is larger. This parallel between the real and the artificial datasets therefore suggests that it is not the difference of $N/S$ ratio between *Dataset1* and *Dataset2* that can explain the different behaviors observed in the periphery of the significant clusters between G-WSPA and ISPA, illustrated on Fig.5 of the paper.

Table 1: Visual comparison of G-WSPA vs ISPA, 9 subjects, 500 data points per subject

| effect size / variance | 0.1 | 0.12 | 0.14 | 0.16 | 0.18 | 0.2 | 0.22 | 0.24 | 0.26 | 0.28 | 0.3 | 0.4 | 0.6 |
|---|---|---|---|---|---|---|---|---|---|---|---|---|---|
| $0.7\pi$ | | | | | | | | | | | | | |
| $0.65\pi$ | | | | | | | | | | | | | |
| $0.6\pi$ | | | | | | | | | | | | | |
| $0.55\pi$ | | | | | | | | | | | | | |
| $0.5\pi$ | | | | | | | | | | | | | |
| $0.45\pi$ | | | | | | | | | | | | | |
| $0.4\pi$ | | | | | | | | | | | | | |
| $0.35\pi$ | | | | | | | | | | | | | |
| $0.3\pi$ | | | | | | | | | | | | | |
| $0.25\pi$ | | | | | | | | | | | | | |
| $0.2\pi$ | | | | | | | | | | | | | |

Table 2: Visual comparison of G-WSPA vs ISPA, 11 subjects, 400 data points per subject

| effect size / variance | 0.1 | 0.12 | 0.14 | 0.16 | 0.18 | 0.2 | 0.22 | 0.24 | 0.26 | 0.28 | 0.3 | 0.4 | 0.6 |
|---|---|---|---|---|---|---|---|---|---|---|---|---|---|
| $0.7\pi$ | | | | | | | | | | | | | |
| $0.65\pi$ | | | | | | | | | | | | | |
| $0.6\pi$ | | | | | | | | | | | | | |
| $0.55\pi$ | | | | | | | | | | | | | |
| $0.5\pi$ | | | | | | | | | | | | | |
| $0.45\pi$ | | | | | | | | | | | | | |
| $0.4\pi$ | | | | | | | | | | | | | |
| $0.35\pi$ | | | | | | | | | | | | | |
| $0.3\pi$ | | | | | | | | | | | | | |
| $0.25\pi$ | | | | | | | | | | | | | |
| $0.2\pi$ | | | | | | | | | | | | | |

Table 3: Visual comparison of G-WSPA vs ISPA, 17 subjects, 250 data points per subject

| effect size / variance | 0.1 | 0.12 | 0.14 | 0.16 | 0.18 | 0.2 | 0.22 | 0.24 | 0.26 | 0.28 | 0.3 | 0.4 | 0.6 |
|---|---|---|---|---|---|---|---|---|---|---|---|---|---|
| $0.7\pi$ | | | | | | | | | | | | | |
| $0.65\pi$ | | | | | | | | | | | | | |
| $0.6\pi$ | | | | | | | | | | | | | |
| $0.55\pi$ | | | | | | | | | | | | | |
| $0.5\pi$ | | | | | | | | | | | | | |
| $0.45\pi$ | | | | | | | | | | | | | |
| $0.4\pi$ | | | | | | | | | | | | | |
| $0.35\pi$ | | | | | | | | | | | | | |
| $0.3\pi$ | | | | | | | | | | | | | |
| $0.25\pi$ | | | | | | | | | | | | | |
| $0.2\pi$ | | | | | | | | | | | | | |

Table 4: Visual comparison of G-WSPA vs ISPA, 21 subjects, 200 data points per subject

| effect size / variance | 0.1 | 0.12 | 0.14 | 0.16 | 0.18 | 0.2 | 0.22 | 0.24 | 0.26 | 0.28 | 0.3 | 0.4 | 0.6 |
|---|---|---|---|---|---|---|---|---|---|---|---|---|---|
| $0.7\pi$ | | | | | | | | | | | | | |
| $0.65\pi$ | | | | | | | | | | | | | |
| $0.6\pi$ | | | | | | | | | | | | | |
| $0.55\pi$ | | | | | | | | | | | | | |
| $0.5\pi$ | | | | | | | | | | | | | |
| $0.45\pi$ | | | | | | | | | | | | | |
| $0.4\pi$ | | | | | | | | | | | | | |
| $0.35\pi$ | | | | | | | | | | | | | |
| $0.3\pi$ | | | | | | | | | | | | | |
| $0.25\pi$ | | | | | | | | | | | | | |
| $0.2\pi$ | | | | | | | | | | | | | |

Table 5: Visual comparison of G-WSPA vs ISPA, 51 subjects, 80 data points per subject

| effect size / variance | 0.1 | 0.12 | 0.14 | 0.16 | 0.18 | 0.2 | 0.22 | 0.24 | 0.26 | 0.28 | 0.3 | 0.4 | 0.6 |
|---|---|---|---|---|---|---|---|---|---|---|---|---|---|
| $0.7\pi$ | | | | | | | | | | | | | |
| $0.65\pi$ | | | | | | | | | | | | | |
| $0.6\pi$ | | | | | | | | | | | | | |
| $0.55\pi$ | | | | | | | | | | | | | |
| $0.5\pi$ | | | | | | | | | | | | | |
| $0.45\pi$ | | | | | | | | | | | | | |
| $0.4\pi$ | | | | | | | | | | | | | |
| $0.35\pi$ | | | | | | | | | | | | | |
| $0.3\pi$ | | | | | | | | | | | | | |
| $0.25\pi$ | | | | | | | | | | | | | |
| $0.2\pi$ | | | | | | | | | | | | | |

Table 6: Visual comparison of G-WSPA vs ISPA, 101 subjects, 40 data points per subject

| effect size / variance | 0.1 | 0.12 | 0.14 | 0.16 | 0.18 | 0.2 | 0.22 | 0.24 | 0.26 | 0.28 | 0.3 | 0.4 | 0.6 |
|---|---|---|---|---|---|---|---|---|---|---|---|---|---|
| $0.7\pi$ | | | | | | | | | | | | | |
| $0.65\pi$ | | | | | | | | | | | | | |
| $0.6\pi$ | | | | | | | | | | | | | |
| $0.55\pi$ | | | | | | | | | | | | | |
| $0.5\pi$ | | | | | | | | | | | | | |
| $0.45\pi$ | | | | | | | | | | | | | |
| $0.4\pi$ | | | | | | | | | | | | | |
| $0.35\pi$ | | | | | | | | | | | | | |
| $0.3\pi$ | | | | | | | | | | | | | |
| $0.25\pi$ | | | | | | | | | | | | | |
| $0.2\pi$ | | | | | | | | | | | | | |

Table 7: Visual comparison of G-WSPA vs ISPA, 201 subjects, 20 data points per subject

| effect size / variance | 0.1 | 0.12 | 0.14 | 0.16 | 0.18 | 0.2 | 0.22 | 0.24 | 0.26 | 0.28 | 0.3 | 0.4 | 0.6 |
|---|---|---|---|---|---|---|---|---|---|---|---|---|---|
| $0.7\pi$ | | | | | | | | | | | | | |
| $0.65\pi$ | | | | | | | | | | | | | |
| $0.6\pi$ | | | | | | | | | | | | | |
| $0.55\pi$ | | | | | | | | | | | | | |
| $0.5\pi$ | | | | | | | | | | | | | |
| $0.45\pi$ | | | | | | | | | | | | | |
| $0.4\pi$ | | | | | | | | | | | | | |
| $0.35\pi$ | | | | | | | | | | | | | |
| $0.3\pi$ | | | | | | | | | | | | | |
| $0.25\pi$ | | | | | | | | | | | | | |
| $0.2\pi$ | | | | | | | | | | | | | |

Table 8: Visual comparison of G-WSPA vs ISPA, 401 subjects, 10 data points per subject

| effect size / variance | 0.1 | 0.12 | 0.14 | 0.16 | 0.18 | 0.2 | 0.22 | 0.24 | 0.26 | 0.28 | 0.3 | 0.4 | 0.6 |
|---|---|---|---|---|---|---|---|---|---|---|---|---|---|
| $0.7\pi$ | | | | | | | | | | | | | |
| $0.65\pi$ | | | | | | | | | | | | | |
| $0.6\pi$ | | | | | | | | | | | | | |
| $0.55\pi$ | | | | | | | | | | | | | |
| $0.5\pi$ | | | | | | | | | | | | | |
| $0.45\pi$ | | | | | | | | | | | | | |
| $0.4\pi$ | | | | | | | | | | | | | |
| $0.35\pi$ | | | | | | | | | | | | | |
| $0.3\pi$ | | | | | | | | | | | | | |
| $0.25\pi$ | | | | | | | | | | | | | |
| $0.2\pi$ | | | | | | | | | | | | | |

#### 2. Influence of the within-subject covariance

**Aim**. Group-level decoding is strongly influenced by the ratio of the inter- and within-subject variance. To study the influence of this ratio, we performed experiments in the paper where the within-subject covariance $\Sigma$ was fixed and the inter-subject variability was parametrically controlled, using our generative model to create a large number of artificial datasets. Here, we study whether our results hold when we change the within-subject variance.

**Experiments**. In this section we vary both the within- and inter-subject variability. We keep the same range of values for $\Theta$, which controls the amount of inter-subject variability:
$\Theta \in \{0.2\pi, 0.25\pi, 0.3\pi, 0.35\pi, 0.4\pi, 0.45\pi, 0.5\pi, 0.55\pi, 0.6\pi, 0.65\pi, 0.7\pi\}$. And we generate five new sets of 14300 datasets using five values for the within-subject covariance matrix:
$\Sigma_1 = \begin{pmatrix} 1 & 0 \\ 0 & 5 \end{pmatrix}, \Sigma_2 = \begin{pmatrix} 3 & 0 \\ 0 & 5 \end{pmatrix}, \Sigma_3 = \begin{pmatrix} 5 & 0 \\ 0 & 5 \end{pmatrix}, \Sigma_4 = \begin{pmatrix} 7 & 0 \\ 0 & 5 \end{pmatrix}, \Sigma_5 = \begin{pmatrix} 9 & 0 \\ 0 & 5 \end{pmatrix}$.
We fix the number of subjects to 21 and generate 200 data points for each subject. Figure 1 illustrates the effect of each of these five covariance matrices on the properties of the generated datasets, for $d = 2$ and $\Theta = 0$. In short, the distinctiveness of the two classes decreases from $\Sigma_1$ to $\Sigma_5$.

**Results**. Our results are shown in Tables 9 to 13. As previously, G-WSPA and ISPA yield more that 50 detections out of the 100 datasets available in each cell in regions that only partially overlap. Both strategies are strongly influenced by the within-subject covariance $\Sigma$, i.e they prove less effective when going from $\Sigma_1$ to $\Sigma_5$, i.e when the within-subject distinctiveness of the patterns decreases. However, the shape of the different regions in this parameter space remains constant: G-WSPA is effective in a rectangle area (blue + green cells), showing that it is not affected by the amount of inter-subject variability; and ISPA is effective in a triangle-like area (blue + red cells). This shows that the qualitative nature of our results does not seem to be affected by the value of $\Sigma$.

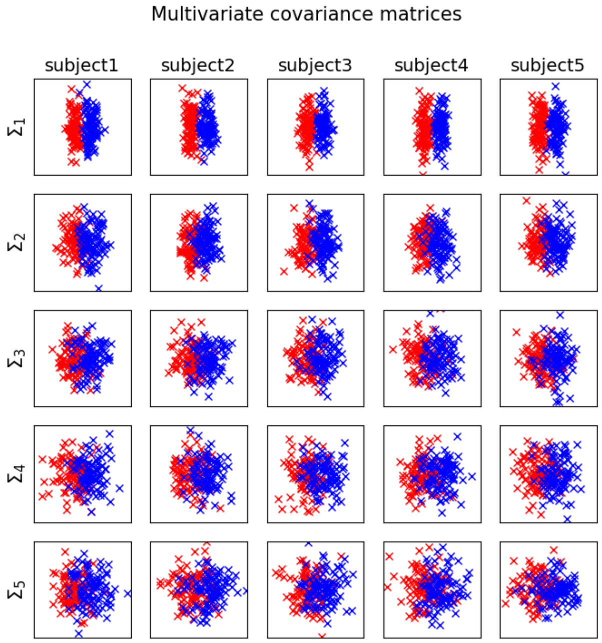


Figure 1: Illustration of the artificial datasets generated with five covariance matrices. Each line is a subpart of a single dataset (5 subjects shown amongst 21), $d = 2$ and $\Theta = 0$.

Table 9: Visual comparison of G-WSPA vs ISPA, $\Sigma_1$

| variance \ effect size | 0.1 | 0.12 | 0.14 | 0.16 | 0.18 | 0.2 | 0.22 | 0.24 | 0.26 | 0.28 | 0.3 | 0.4 | 0.6 |
|---|---|---|---|---|---|---|---|---|---|---|---|---|---|
| $0.7\pi$ | | | | | | | | | | | | | |
| $0.65\pi$ | | | | | | | | | | | | | |
| $0.6\pi$ | | | | | | | | | | | | | |
| $0.55\pi$ | | | | | | | | | | | | | |
| $0.5\pi$ | | | | | | | | | | | | | |
| $0.45\pi$ | | | | | | | | | | | | | |
| $0.4\pi$ | | | | | | | | | | | | | |
| $0.35\pi$ | | | | | | | | | | | | | |
| $0.3\pi$ | | | | | | | | | | | | | |
| $0.25\pi$ | | | | | | | | | | | | | |
| $0.2\pi$ | | | | | | | | | | | | | |

Table 10: Visual comparison of G-WSPA vs ISPA, $\Sigma_2$

| variance \ effect size | 0.1 | 0.12 | 0.14 | 0.16 | 0.18 | 0.2 | 0.22 | 0.24 | 0.26 | 0.28 | 0.3 | 0.4 | 0.6 |
|---|---|---|---|---|---|---|---|---|---|---|---|---|---|
| $0.7\pi$ | | | | | | | | | | | | | |
| $0.65\pi$ | | | | | | | | | | | | | |
| $0.6\pi$ | | | | | | | | | | | | | |
| $0.55\pi$ | | | | | | | | | | | | | |
| $0.5\pi$ | | | | | | | | | | | | | |
| $0.45\pi$ | | | | | | | | | | | | | |
| $0.4\pi$ | | | | | | | | | | | | | |
| $0.35\pi$ | | | | | | | | | | | | | |
| $0.3\pi$ | | | | | | | | | | | | | |
| $0.25\pi$ | | | | | | | | | | | | | |
| $0.2\pi$ | | | | | | | | | | | | | |

Table 11: Visual comparison of G-WSPA vs ISPA, $\Sigma_3$

| variance \ effect size | 0.1 | 0.12 | 0.14 | 0.16 | 0.18 | 0.2 | 0.22 | 0.24 | 0.26 | 0.28 | 0.3 | 0.4 | 0.6 |
|---|---|---|---|---|---|---|---|---|---|---|---|---|---|
| $0.7\pi$ | | | | | | | | | | | | | |
| $0.65\pi$ | | | | | | | | | | | | | |
| $0.6\pi$ | | | | | | | | | | | | | |
| $0.55\pi$ | | | | | | | | | | | | | |
| $0.5\pi$ | | | | | | | | | | | | | |
| $0.45\pi$ | | | | | | | | | | | | | |
| $0.4\pi$ | | | | | | | | | | | | | |
| $0.35\pi$ | | | | | | | | | | | | | |
| $0.3\pi$ | | | | | | | | | | | | | |
| $0.25\pi$ | | | | | | | | | | | | | |
| $0.2\pi$ | | | | | | | | | | | | | |

Table 12: Visual comparison of G-WSPA vs ISPA, $\Sigma_4$

| variance \ effect size | 0.1 | 0.12 | 0.14 | 0.16 | 0.18 | 0.2 | 0.22 | 0.24 | 0.26 | 0.28 | 0.3 | 0.4 | 0.6 |
|---|---|---|---|---|---|---|---|---|---|---|---|---|---|
| $0.7\pi$ | | | | | | | | | | | | | |
| $0.65\pi$ | | | | | | | | | | | | | |
| $0.6\pi$ | | | | | | | | | | | | | |
| $0.55\pi$ | | | | | | | | | | | | | |
| $0.5\pi$ | | | | | | | | | | | | | |
| $0.45\pi$ | | | | | | | | | | | | | |
| $0.4\pi$ | | | | | | | | | | | | | |
| $0.35\pi$ | | | | | | | | | | | | | |
| $0.3\pi$ | | | | | | | | | | | | | |
| $0.25\pi$ | | | | | | | | | | | | | |
| $0.2\pi$ | | | | | | | | | | | | | |

Table 13: Visual comparison of G-WSPA vs ISPA, $\Sigma_5$

| variance \ effect size | 0.1 | 0.12 | 0.14 | 0.16 | 0.18 | 0.2 | 0.22 | 0.24 | 0.26 | 0.28 | 0.3 | 0.4 | 0.6 |
|---|---|---|---|---|---|---|---|---|---|---|---|---|---|
| $0.7\pi$ | | | | | | | | | | | | | |
| $0.65\pi$ | | | | | | | | | | | | | |
| $0.6\pi$ | | | | | | | | | | | | | |
| $0.55\pi$ | | | | | | | | | | | | | |
| $0.5\pi$ | | | | | | | | | | | | | |
| $0.45\pi$ | | | | | | | | | | | | | |
| $0.4\pi$ | | | | | | | | | | | | | |
| $0.35\pi$ | | | | | | | | | | | | | |
| $0.3\pi$ | | | | | | | | | | | | | |
| $0.25\pi$ | | | | | | | | | | | | | |
| $0.2\pi$ | | | | | | | | | | | | | |

#### 3. Influence of spatial smoothing on the results obtained on the real fMRI datasets

**Aim**. The preprocessing steps performed in the paper on the two fMRI datasets did not include any spatial smoothing. Since there is a debate in the literature on the validity and interest of such smoothing when performing MVPA, we here study the influence of the smoothing on our results.

**Experiments**. We replicate the same experiments as in the paper by adding some spatial smoothing on the beta maps that are used to construct the inputs of the classifier. We use the Gaussian kernel implemented in SPM to perform the smoothing, with full-width at half-maximum values of $3mm$ and $6mm$.

**Results**. For clarity, we denote as *Dataset1-s3* and *Dataset2-s3* the versions of the two datasets with the $3mm$ smoothing, and *Dataset1-s6* and *Dataset2-s6* with the $6mm$ smoothing. The results are presented in the same way as in the paper for the unsmoothed data: Figures. 2 and 3 show the thresholded statistical maps, displaying the significant clusters; Figures. 4 and 5 present the analysis of the size of the main cluster and its peak statistic value, for each dataset, each hemisphere and each size of the searchlight radius; finally, Figures. 6 and 7 describe how the main clusters obtained with G-WSPA and ISPA overlap.

Overall, the results obtained with smoothing are consistent with what we found with the unsmoothed data: clusters were found in the same regions of the brain, their size increased with the value of the searchlight radius and the patterns of overlap between the clusters found by G-WSPA and ISPA were overall similar. One notable difference is the fact that the size of the activated clusters found by ISPA for *Dataset2-s6* is as large as the ones found by G-WSPA, which is not the case with smaller or no smoothing (see bottom-left graph in Figure. 5, compared with the one in Figure. 4 or with Figure. 4 in the paper). Congruently, the pattern of overlapping of the clusters found by the two strategies is modified for *Dataset2-s6* when compared with the one found with smaller or no smoothing (see the right graph of Figure. 7, compared to the right graph on Figure. 6 and Figure. 6 in the paper): the number of voxels detected only by G-WSPA (green voxels) decreased in an important proportion. This is consistent with the putative explanation that was put forward in the paper, that stated that the large amount of green voxels found in *Dataset2* could be caused by a large inter-individual variability: indeed, one of the effect of spatial smoothing is often to reduce this inter-individual variability.

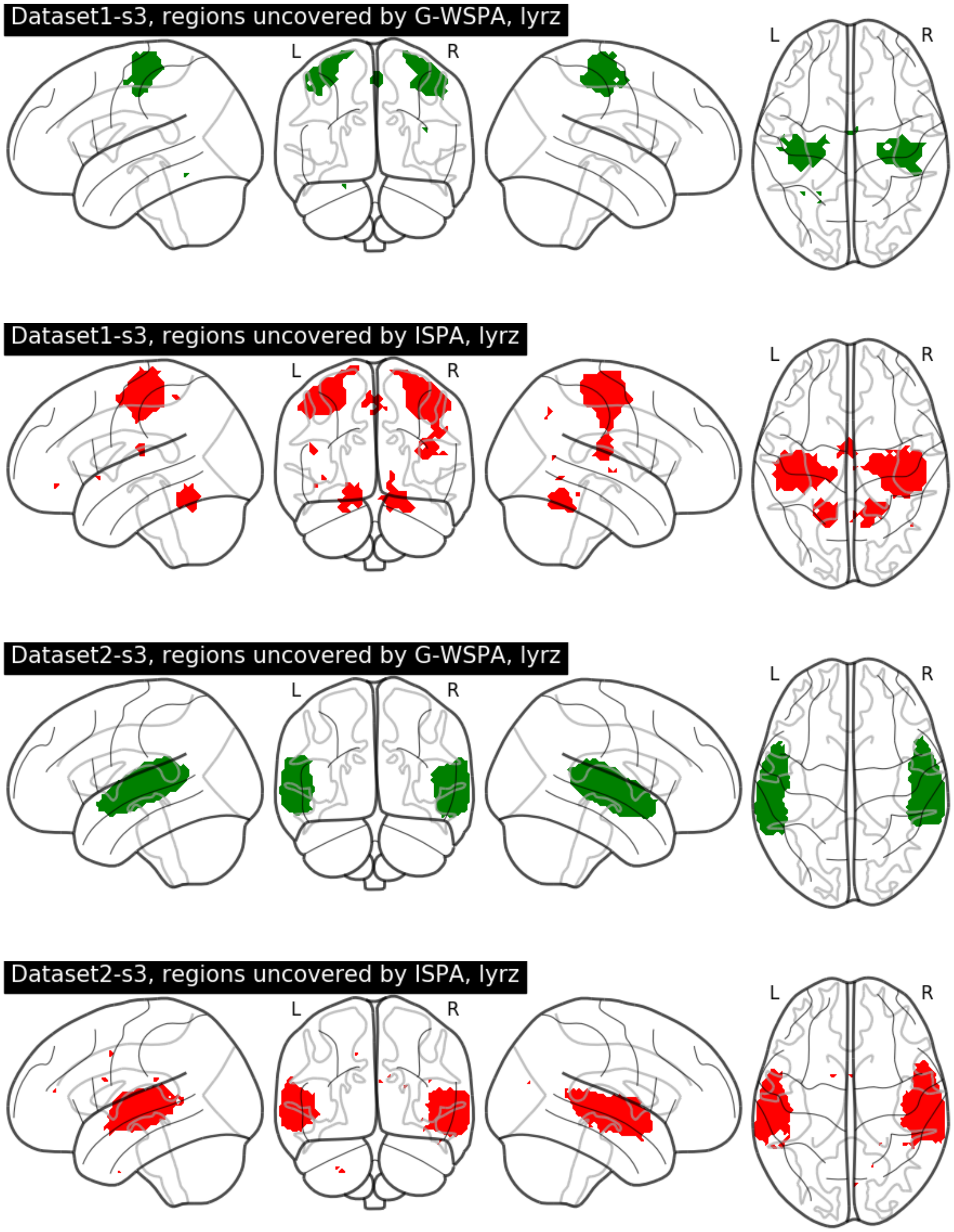


Figure 2: Illustration of the results of the group-level searchlight decoding analysis for a $6mm$ radius with *Dataset1-s3* and *Dataset2-s3*. Top two rows: *Dataset1-s3*; bottom two rows: *Dataset2-s3*. Brain regions found significant using G-WSPA and ISPA are respectively depicted in green and red.

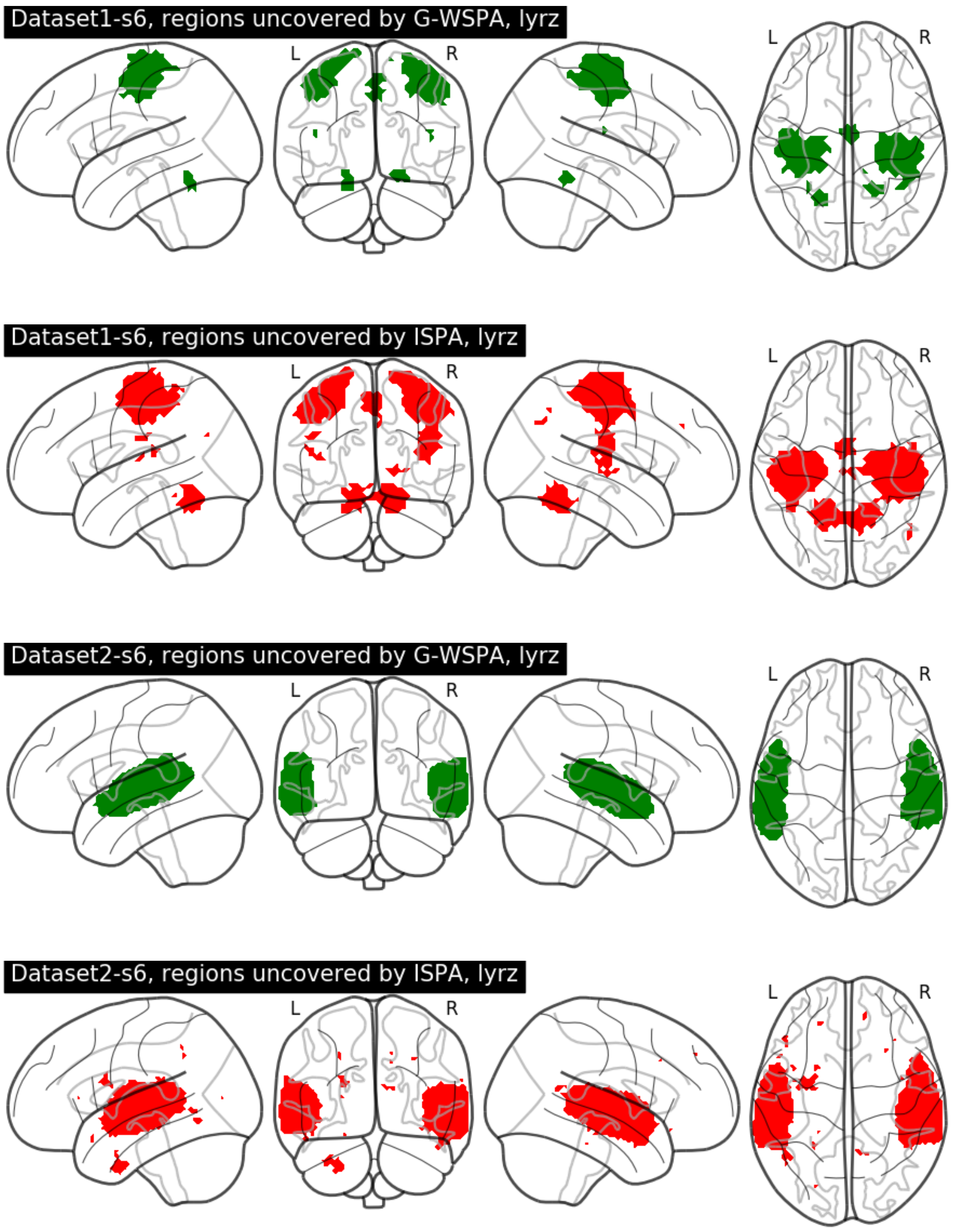


Figure 3: Illustration of the results of the group-level searchlight decoding analysis for a 6*mm* radius with *Dataset1-s6* and *Dataset2-s6*. Top two rows: *Dataset1-s6*; bottom two rows: *Dataset2-s6*. Brain regions found significant using G-WSPA and ISPA are respectively depicted in green and red.

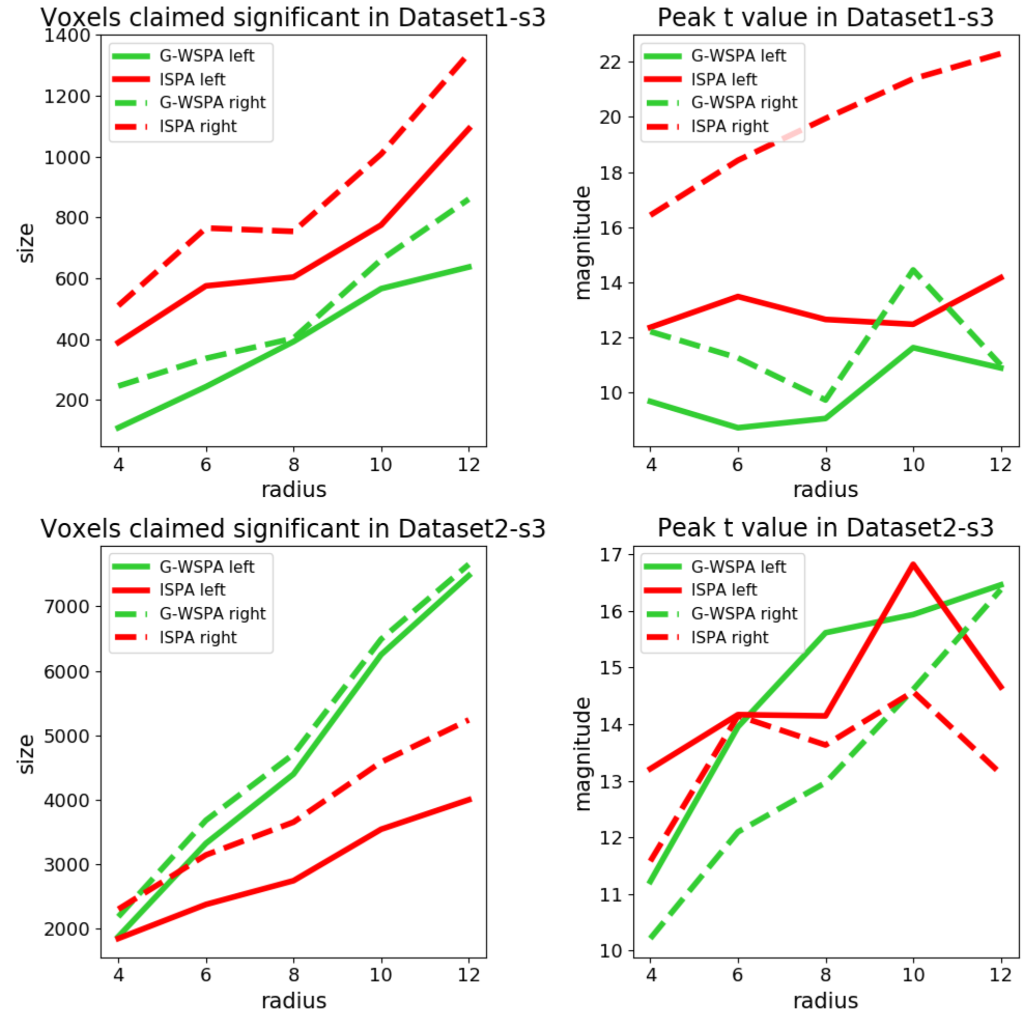


Figure 4: Quantitative evaluation of the results obtained on the smoothed fMRI datasets for G-WSPA (green curves) and ISPA (red curves), $3mm$ Gaussian kernel. Solid and dashed lines for the largest cluster respectively in the left and right hemispheres. Left column: size of the significant clusters. Right column: peak $t$ statistic. Top vs. bottom row: results for *Dataset1-s3* and *Dataset2-s3* respectively.

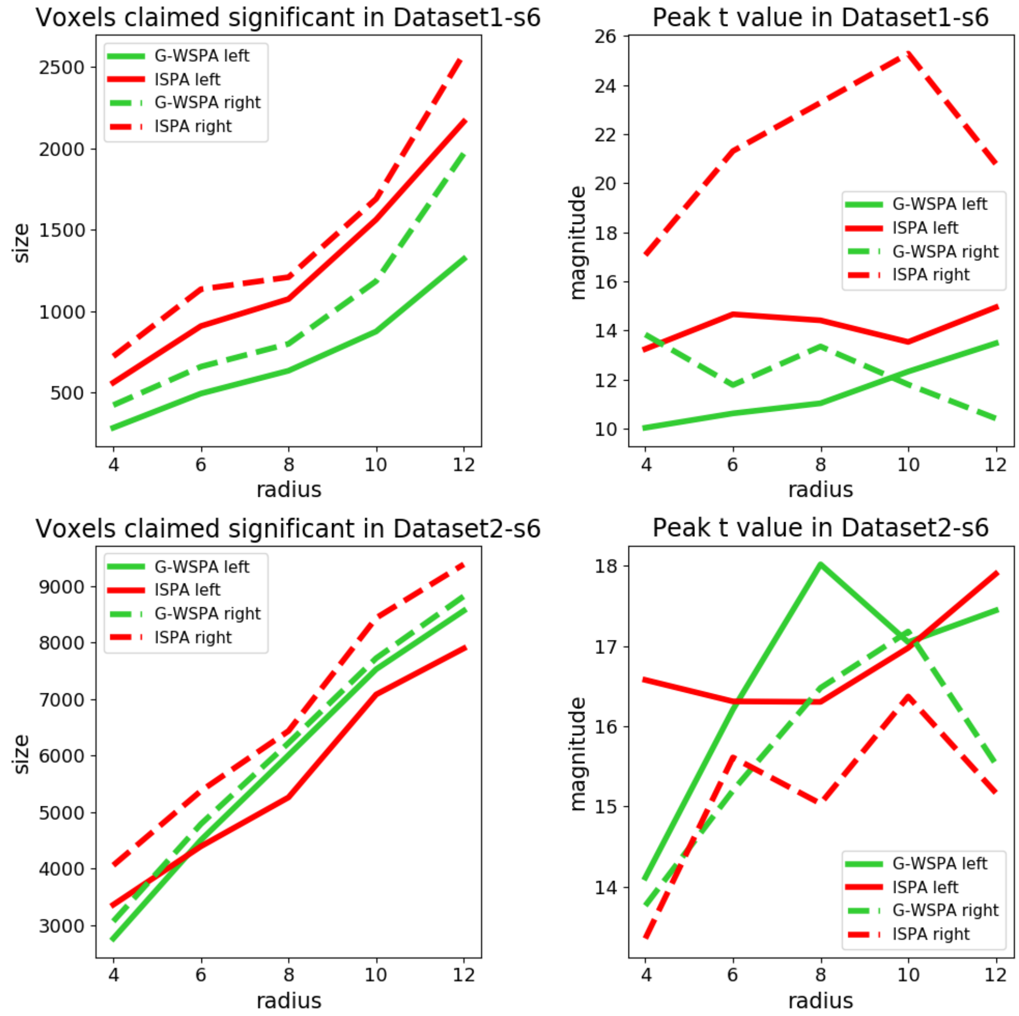


Figure 5: Quantitative evaluation of the results obtained on the smoothed fMRI datasets for G-WSPA (green curves) and ISPA (red curves), $6mm$ Gaussian kernel. Solid and dashed lines for the largest cluster respectively in the left and right hemispheres. Left column: size of the significant clusters. Right column: peak $t$ statistic. Top vs. bottom row: results for *Dataset1-s6* and *Dataset2-s6* respectively.

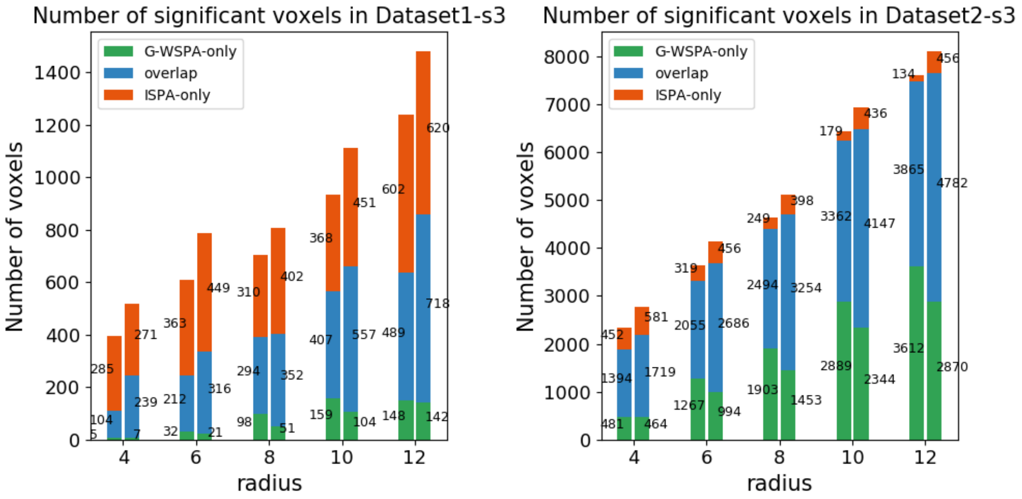


Figure 6: Comparison of the voxel counts detected by G-WSPA-only (green), ISPA-only (red) or both (blue) for the different values of the searchlight radius, in *Dataset1-s3* (left) and *Dataset2-s3* (right)

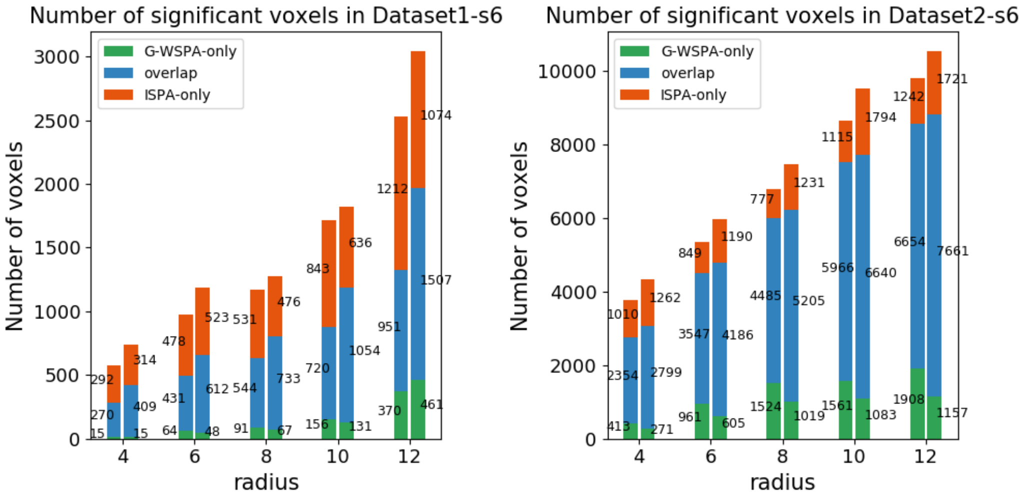


Figure 7: Comparison of the voxel counts detected by G-WSPA-only (green), ISPA-only (red) or both (blue) for the different values of the searchlight radius, in *Dataset1-s6* (left) and *Dataset2-s6* (right).
